# Recasting adaptation as strategy inference

**DOI:** 10.1101/2025.03.24.645064

**Authors:** Sami Beaumont, Mehdi Khamassi, Philippe Domenech

## Abstract

Flexible adaptation to uncertain and changing environments requires dynamic adjustments in behavioral strategies. While classical learning theories emphasize incremental strengthening of local stimulus-action associations in adaptation, emerging evidence suggests that global-level strategy representations may enable rapid inference of adaptive behaviors, thus promoting efficient decision-making. However, it remains unclear to what extent direct inference over putative strategies can fully account for human adaptation across diverse statistical contexts. Here, we demonstrate clear behavioral markers supporting the broad use of inference over strategies in human adapting to rapid changes. These markers are fully explained solely by a novel model of inference over a structured space of strategies. We further show that inference over strategies is influenced by latent contextual statistics that are beyond the scope of models based on incremental learning. Taken together, these results establish the importance of direct inference over an abstract strategy space for flexible adaptation in humans.

---

Flexible adaptation to unexpected changes is a hallmark of effective decisionmaking, often manifested as rapid behavioral switches [1, 2, 3, 4, 5, 6]. These switches are pervasive and primarily driven by efficiency, balancing the exploration-exploitation trade-off [7], but may also reflect idiosyncrasies specific to individuals or species [8]. These rapid behavioral switches have been formalized as *strategy changes*, i.e. at an integrated, global and abstract level of representation of all available stimulus-actions associations [7, 9, 10, 11, 12]. Hence, a *strategy* is a mapping between stimuli and actions that forms a cohesive policy consistently guiding an individual choices across a series of successive trials. Extensive experimental data support the key role of this level of representation for adaptive behavior in rodents [2, 3, 13], non-human primates [4, 6, 14, 15, 16] and humans [5, 7, 11, 17].

The extent to which inference over a collection of putative strategies can fully account for human adaptation across diverse statistical contexts remains an open question. Environmental feedback can induce incremental modifications of a strategy [5, 9, 12], typically by changing the associative strength of a single stimulus-action pair while keeping that of the other pairs unchanged. However, a growing body of evidence suggests that behavioral adaptability in humans can also be decoupled from such gradual feedback integration : as seen in one-shot learning [18], instruction sensitivity [19, 20, 21] or generalization from previous experience to different contexts [15, 22]. In this work, we explore the idea that humans can represent strategies at an abstract cognitive level within a structured strategy space, and rapidly adapt to changes by flexibly navigating putative task-relevant strategies (see Figure 1A). Such mechanism may dominate over more incremental adaptation processes, resulting in behavioral adjustments that are highly malleable and significantly influenced by latent environmental statistics.

**Figure 1:**
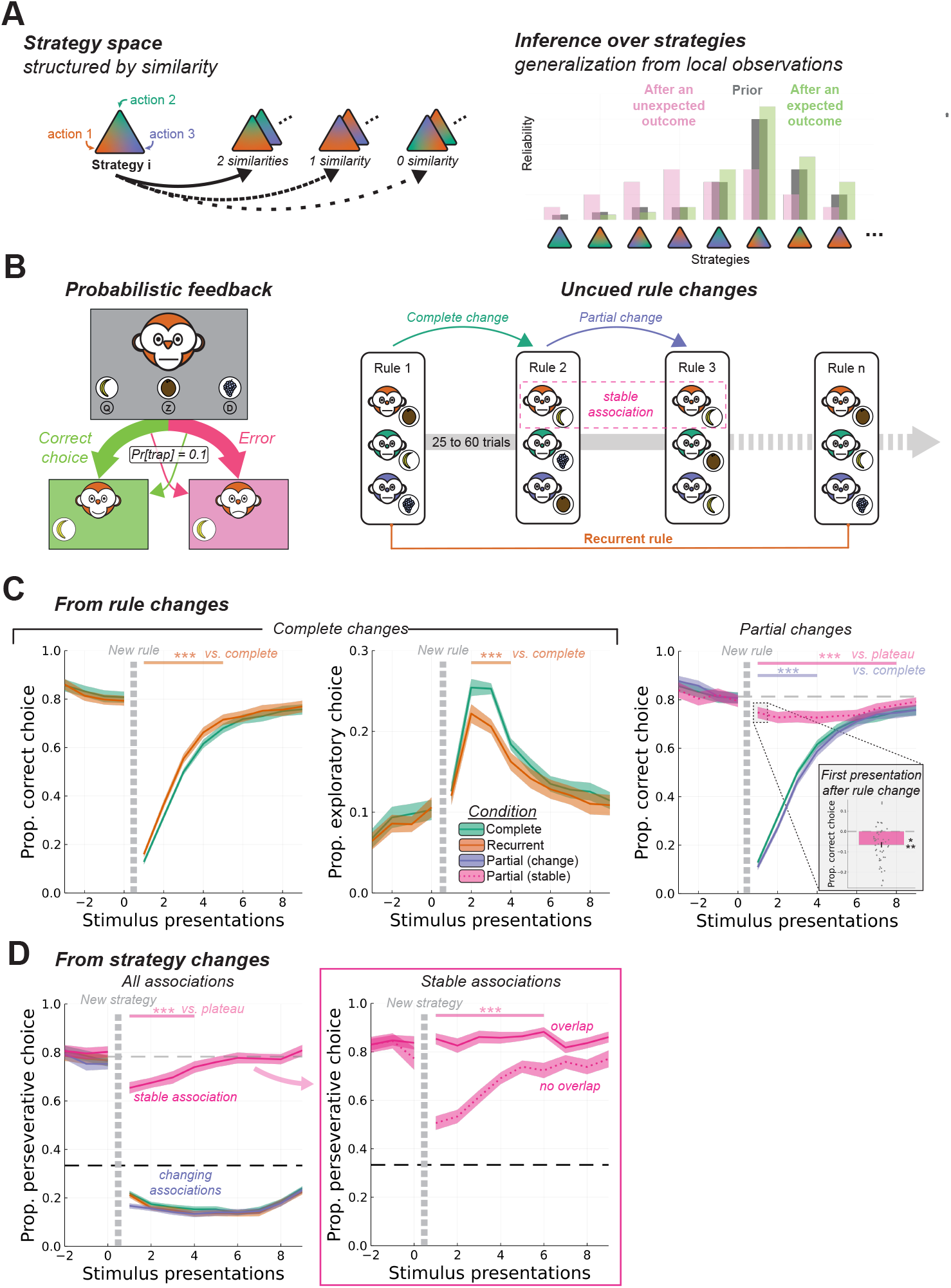
Theoretical framework and experimental task. **A:** Schematic of the theoretical framework and hypotheses. Example of global strategies related by similarity (left panel). Inference over the space of strategies, with similar strategies having close reliabilities (right panel). After the observation of an expected outcome (i.e. in line with the prior), the reliability distribution narrows around the same mode (green). After an unexpected outcome, uncertainty increases and the reliability distribution shifts, affecting several similar strategies (pink). **B:** The Monkey Feeding Task. On each trial, participants were presented one of the three monkeys, and had to choose one of the three fruits. Once they made a choice, a feedback screen indicated the monkey, the chosen fruit and the positive or negative reaction of the monkey. Crucially, only 90% of feedbacks reflected the actual rule. Rules changes occurred without notice ten trials after reaching a performance criterion, and could be either complete or partial (i.e. one association stayed the same). In the former case, the next rule could be the recurrent one. **C:** Behavioral performance locked on rule changes (mean ± standard error). The x-axis represents successive presentations of each stimulus (and not consecutive trials). Ribbons on the top represent statistically significant effects after correcting for multiple comparisons with a cluster based permutation test. Left: After a change to the recurrent rule, performance increased significantly faster compared to changes for new rules (***: p < 0.001). Middle: After a change to the recurrent rule, non perseverative errors (i.e. exploratory responses) were significantly reduced compared to changes for new rules. Right: After a partial change of rules, performance increased significantly more slowly than after a complete change, and the performance for the stable association significantly decreased. Insert : Difference between the performance in the first presentation of the stable association after the rule change and the performance during the plateau (population mean ± standard error). **D:** Behavioral performance locked on strategy changes as detected by the HMM (dashed line: chance level). Left: For all associations, the average proportion of perseverative responses abruptly dropped after a strategy change. Right: For stable associations, the average proportion of perseverative responses decreased only after a non overlapping strategy change.

Typical experimental paradigms used to study adaptation processes in changing environments cannot separate incremental learning from direct inference at the strategy level for several reasons. First, global strategies must be distinguishable from local actions. This requires studying behavior in environments with more than one stimulus, or state, each associated with different possible choices, which is generally not the case in probabilistic reversal or bandit tasks [23]. Even in tasks requiring more complex behavioral strategies, deducible relationship between associations (e.g. mutual exclusion) often facilitate updating associations with unseen stimuli from a partial observation, thereby blurring the contribution of inference at the global level (for example in [5, 24]). Second, although some see abrupt behavioral switches as evidence against incremental learning [1], it is possible for learning parameters (such as learning rate or inverse temperature) to transiently assume extreme values, mimicking discrete strategic switching effects [25, 26, 27, 28].

To study the contribution of inference over strategies to human adaptation, we designed an original task addressing these challenges. Our study reveals several clear-cut behavioral signatures of inference over strategies when human adapt to rapid changes that could only be comprehensively explained by a novel model of inference over a structured space of strategies, not by incremental learning mechanisms alone. We further show that inference over strategies is influenced by latent statistics specific to the environment structure (which we refer to as contexts) that are beyond the scope of models based on incremental learning.

We administered a modified Wisconsin Card Sorting Task (Figure 1B) to two independent groups of healthy human participants (main sample, n=51; additional sample, n=54). Participants were asked to find the preferred fruit (banana, coconut or grapes) for each of 3 monkeys, distinguished by different coat colors. In each trial, they were presented with one monkey and asked to choose one of the three fruits. Then, they received feedback indicating whether the monkey was satisfied with their choice. Importantly, participants were only instructed that each monkey had a single preferred fruit at any given time, that this preference could change overtime, and that a given fruit could be preferred by one or two monkeys simultaneously. This design prevented the use of heuristics and deductive reasoning to infer correct associations. In this task, the goal was to maximize positive feedback, with participants receiving a monetary bonus proportional to their performance. Feedback was unreliable (meaning it did not match the underlying rule of association between fruit and monkey) in 10% of the trials. Hence, participants had to infer the underlying rule by trial and errors over successive trials, while these rules changed periodically without explicit cues. We will henceforth distinguish between “*rule changes*”, when referring to actual changes in correct fruit-monkey associations, and “*strategy changes*” when referring to changes in behavioral strategies observed in participants’ responses.

Participants completed several sessions of the task over 15 days. Unbeknownst to them, the proportion and the type of rule changes were systematically manipulated to create distinct hidden structures of the environment - referred to as experimental “*contexts*”. This design allowed us to test whether participants would leverage such underlying structure to efficiently infer global strategies from local observations. Rule changes could either be complete (*all monkey-fruit associations changed*) or partial (*one or two associations remained stable*; see Figure 1B, right panel). More precisely, participants in the main sample (n=51) performed the task in two different experimental contexts, the order of which was counterbalanced across the four experimental sessions: (1) in one context rule changes could be either complete or partial (with one stable association; “*partial vs complete changes*”), (2) in the other context rule changes were always partial with one or two stable associations (“*only partial changes*”). We also administered the task to an additional sample of participants (n=54) in three additional contexts (three sessions in counterbalanced order): one with complete rule changes only, one with a mix of complete and partial rule changes (with one stable association), and one with partial rule changes only (with one or two stable associations). This additional sample was collected to replicate and extend key behavioral markers observed in the main sample and to optimize experimental environments to better disentangle potential confounds (for example by tightly controlling for apparent local stimulus-response volatility). The characteristics of all the experimental contexts (for both main and additional sample) are detailed in Supplementary Figure S1 (see also Material and Methods). By default, all results reported afterward pertain to the main sample performing the task in the “*partial vs complete changes*” (PVC) or the “*only partial changes*” (OPC) contexts, unless specifically stated otherwise.

## Behavioral markers of inference over strategies

### A working memory buffer at the strategic level explains observed changes after a recurrent rule

In the “*partial vs complete changes*” (PVC) context, all monkeys simultaneously modified their preferred fruit simultaneously during complete rule changes (see Figure 1B, green arrow, 26 out of the 36 rule changes per session in the main sample). The average proportion of correct responses initially dropped upon encountering each stimulus for the first time but gradually increased as participants inferred the new underlying rule (Figure 1C, left panel). Due to the task design, with three possible choices, incorrect responses could be classified as either (1) perseverative errors, where the participants erroneously chose the action that was correct under the previous rule, or (2) non perseverative errors, corresponding to purely exploratory choices, where participants selected an action that was neither correct under the previous nor the current rule.

Complete changes could lead to either a new rule (16 changes) or a previously encountered “recurrent” rule (10 changes). This single recurrent rule allowed us to test whether rule memories exist at the global level of strategies, as opposed to the local level of individual stimulus-action pairs. Indeed, the rapid retrieval of previously used strategies from a working memory buffer is a hallmark of inference mechanisms at the global (strategic) level—an effect not predicted by classical incremental learning models. In line with the strategy inference hypothesis, participants identified correct responses more quickly after a change to the recurrent rule (Figure 1 panel C, left and middle). We tested the difference of correct responses between the new rule and the recurrent rule condition over a 10-stimulus presentation window following a rule change (approximately 30 trials), correcting for multiple comparisons with cluster-based permutation (significant cluster from stimulus presentation 1 to 5, corrected *p* < 0.001). Interestingly, participants also explored less when presented with the recurrent rule than with a new rule (Figure 1C middle panel; significant cluster from stimulus presentation 2 to 4, corrected *p* < 0.001).

However, because only a single rule recurred throughout an entire experimental session of the PVC context, this effect could also be driven by prior beliefs about the frequency of local associations biasing exploration after a rule change rather than reflecting the retrieval of a strategy memory. To test whether the memory effect was specific to the repetition of a global rule rather than the increased frequency of local associative links, we compared choice behavior after the change toward rules composed of rare monkey-fruit associations (i.e., associations not used in at least the previous 5 rules) with rules composed of more frequent associations. Strikingly, correct and exploratory response patterns for frequent and rare monkey-fruit associations completely overlapped (Supplementary Figure S2; no significant cluster survived the cluster-based permutation correction), demonstrating that the observed adaptation to recurrent rules was not driven by the observed frequency of local stimulus-action associations. Finally, we replicated the effect of recurrence on adaptation in the additional sample of participant using a context including only complete rule changes (“*only complete changes*” or OCC context), where three recurrent rules pseudo-randomly alternated with new rules (Supplementary Figure S3, left panel; significant cluster from presentations 5 to 10 after rule change, corrected *p* < 0.001). Taken together, these results are consistent with previous studies showing accelerated adaptation to rule changes towards previously encountered strategies [5, 29] and extend this effect to open contexts mixing recurrent and new rules.

### Adaptation to partial rule changes reveals generalization from local stimulus-response-outcome observations to global strategies

In the “*partial vs complete changes*” (PVC), partial rule changes occurred 10 out of 36 times, modifying two monkey-fruit associations while keeping one unchanged (referred to as the “*stable association*”, see Figure 1B, blue arrow). We reasoned that if participants used inference at the global level of strategies, a few observations could generalize to the whole strategy, leading to faster adaptation but potentially impairing the performance of the stable association. Confirming this prediction, the proportion of correct responses for the stable association transiently decreased after a partial rule change (compared to the pre-change performance plateau, from presentation 1 to 8; *p* < 0.001; Figure 1C right panel). Strikingly, this interference was already measurable from the very first presentation of the stable stimulus after the rule change, as shown by a comparison of performance on this trial with performance at plateau (Figure 1C insert; median performance drop from plateau : −0.051, *CI*_95_ = [−0.090, −0.038], Wilcoxon signed rank test *Z* = 182.5, *p* < 0.001). Moreover, this effect was replicated in two other contexts, in the task administered to the independent group of participants, for partial rule changes with either one or two stable associations (median performance drop from plateau with one stable association in mixed complete/partial changes context: 0.019, *CI*_95_ = [− 0.053, −0.004], Wilcoxon signed rank test *Z* = 481, *p* < 0.025; in only partial changes context : −0.033, *CI*_95_ = [−0.057, −0.011], Wilcoxon signed rank test *Z* = 421, *p* = 0.006; with two stable associations : 0.028, *CI*_95_ = [−0.044, −0.013], Wilcoxon signed rank test *Z* = 366, *p* = 0.001; Supplementary Figure S3 insert).

The existence of such an interference effect is a direct prediction of inference over strategies, since from an incremental learning perspective without global adaptive parameters – which assumes that local adaptation occurs independently for each monkey-fruit association – no change should be observable for the stable association after the rule change in these variants of our task (as an illustration, see the simulation of a standard Q-learning model shown in Supplementary Figure S4 top left corner).

In addition, while simple incremental learning would predict a similar learning speed for new associations, whether the other changed or not, the detection of change at the strategic level would be delayed after partial rule changes, leading to slower relearning compared to complete rule changes. Consistent with the strategy inference hypothesis, partial rule changes led to slower increasing of correct responses compared to complete rule changes in the same context, in participants of the main sample (significant cluster from stimulus presentations 1 to 4; corrected *p* < 0.001), though this effect was not replicated in the additional sample.

### Recovered strategy changes in participants reveal dynamic and adaptive patterns at the strategy level

Assuming that adaptation to a rule change occurs as a global strategy change, and not a local adjustment of associative weights, we sought to detect such switches in participants’ behavior. To this end, we used a hidden Markov model (HMM) designed to infer the most likely position and identity of each strategy change based on the participants’ choice patterns (see Material and Methods). Inference on the HMM yielded the most likely strategy used on each trial, from among the 27 possible combinations (3^3^ = 27 possible monkey-fruit mappings in our task) and a 28^*th*^ strategy corresponding to unbiased random choices for every stimulus. Here again, a strategy corresponds to a combination of monkey-fruit associations found by the HMM to best account for the participant’s current behavior, while a rule corresponds to the currently correct set of monkey-fruit associations from the point of view of the task. Next, we classified the first strategy change after a rule change according to the degree of overlap between the new strategy and the previous one (i.e. the number of similar stimulus-action associations), with overlapping changes comprising strategies with 1 or 2 similarities, and non-overlapping changes strategies with 0 similarities or the random strategy (Figure 2A). Note that the main goal of this analysis was to locate more precisely the timings of strategy changes following rule changes. Constraining the space of latent states to the set of possible complementary strategies in the task thus provided greater sensitivity for detecting strategy changes.

**Figure 2:**
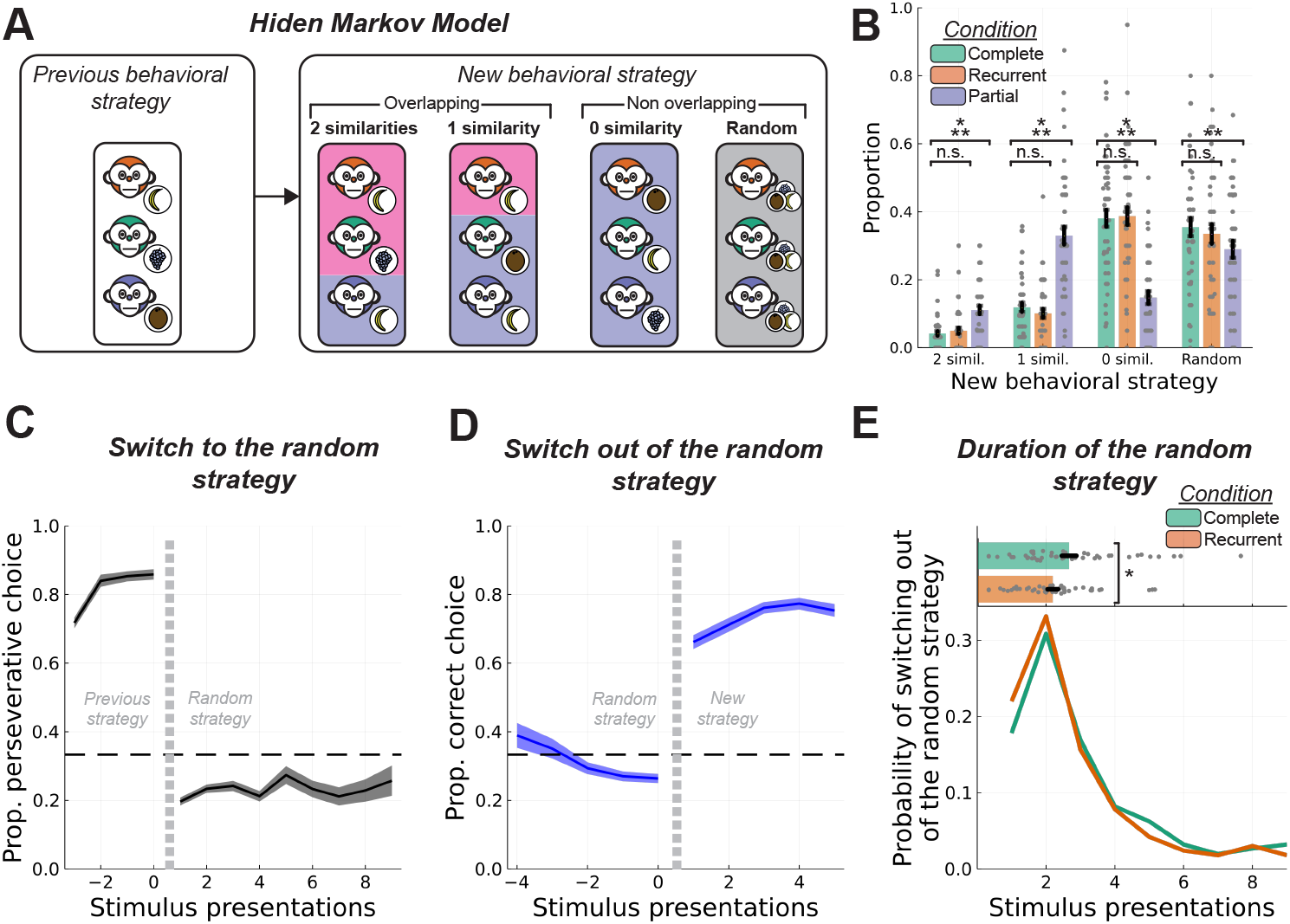
Types and dynamics of strategy changes. **A:** After a rule change, participants’ behavioral strategies changed either for an overlapping strategy (with one or two similar associations), or for a non-overlapping strategy. The later could be the random strategy. **B:** Distribution for strategy changes across experimental conditions (population mean± standard error). Overlapping strategy changes were over-represented after partial rule changes, while 0-similarity strategy changes were over-represented after complete rule changes. The random strategy was used almost as frequently regardless of the type of rule change, although slightly more frequently after a complete rule change. **C:** As with the other strategy changes, perseverative choices decreased sharply when switching to the random strategy. **D:** Switching out of the random strategy was characterized by a sudden increase of correct choices rather than a progressive learning curve. **E:** Duration of the random strategy after complete rule changes (new or recurrent rule). After a change to a recurrent rule, the random strategy was used for fewer trials (2.198 ± 0.162 stimulus presentations) than after a new rule (2.672 ± 0.220 stimulus presentations). ***: p < 0.001, **: p < 0.01, *: p < 0.05, n.s.: non significant.

Perseverative responses locked on strategy changes revealed abrupt behavioral transitions (Figure 1D), thus validating our HMM to accurately detect strategy changes. For changing associations, perseverative responses abruptly dropped around strategy changes (Figure 1D left panel). Remarkably, the decrease of performance for stable associations in partial rule changes (interference effect) was entirely explained by unwarranted non overlapping strategy changes (Figure 1D right panel).

Moreover, strategy changes inferred from the behavior of the participants reflected the conditions in the task (Figure 2B) : 0 similarity strategy changes were more frequent after complete rule changes than after partial rule changes (complete : 0.381 ± 0.025, partial : 0.147 *±* 0.019, Wilcoxon signed rank test *Z* = 1210, *p* < 0.001) and 1 similarity strategy changes were more frequent after partial rule changes than after complete rule changes (complete : 0.119 *±* 0.013, partial : 0.330 *±* 0.026, Wilcoxon signed rank test *Z* = 18, *p* < 0.001). Moreover, 2 similarities strategy changes were overall rare, matching with the absence of corresponding rule changes in this context, but significantly more frequent after partial rule changes than after complete rule changes (complete : 0.042 *±* 0.007, partial : 0.111 *±* 0.012, Wilcoxon signed rank test *Z* = 126, *p* < 0.001). Interestingly, random strategy changes were frequent in all conditions (although slightly less used in the partial rule change condition (complete : 0.35 ± 0.027, partial : 0.29*±* 0.025, Wilcoxon signed rank test *Z* = 894, *p* = 0.005)).

To further test our hypothesis that partial rule changes hindered inference at the strategy level, we computed the latency from actual rule changes to inferred behavioral strategy changes as the average number of stimulus presentations inbetween. Using the number of presentations, and not trials, allowed to cancel out the effect of having a variable number of changing associations (2 or 3). The latency was significantly higher after partial rule changes (3.07 *±* 0.19 presentations) than after complete new rule changes (2.33 ± 0.18 presentations) for all strategy changes excepted 2 similarities strategy changes – which were the least used strategy changes, leading to high variability of latencies (all strategy changes : Wilcoxon signed rank test *Z* = 180, *p* < 0.001; 0 similarity: *Z* = 217, *p* = 0.009; 1 similarity : *Z* = 111, *p* < 0.001; 2 similarities : *Z* = 162.5, *p* = 0.152; random : *Z* = 153, *p* < 0.001). No significant differences were found when comparing the latency after recurrent (2.06 ± 0.14 presentations) and new rule changes (all strategy changes : *Z* = 784, *p* = 0.16; 0 similarity : *Z* = 777, *p* = 0.053; 1 similarity : *Z* = 337, *p* < 0.833; 2 similarities: *Z* = 121, *p* = 0.563; random : *Z* = 405, *p* = 0.095; see Supplementary Figure S5).

As with other strategy changes, transitions into and out of the random strategy were abrupt (Figure 2C & D). However, even though this strategy was designed as a fully random one in the HMM (see Material and Methods), choice patterns classified within the random strategy displayed a slight bias away from perseverative choices (Figure 2C). In addition, participants used this strategy for fewer trials when the true rule was the recurrent one (Figure 2C insert; median duration difference between recurrent and new : −0.211, *CI*_95_ = [−0.666, −0.025] Wilcoxon signed rank test *Z* = 324, *p* = 0.029). Overall, these results indicate that bouts of behavior classified as random were likely an active and directed exploration strategy aimed at gathering evidence when the correct strategy could not be identified, rather than unspecific selection noise.

### Strategy-level inference models provide a more comprehensive and parsimonious account of behavior than incremental learning models

Although the behavioral results demonstrate global effects, these can be the consequence of either direct inference of strategies, or global adaptive control of learning parameters in incremental learning models [28, 30]. In order to separate inference over strategies from incremental learning, we assessed three broad families of generative models (see Material and Methods and Figure 3A). The first one consisted of adaptive Q-learning models, with dynamically adjusted learning rate, inverse temperature or both. This Q-learning family is based entirely on incremental learning, and global effects can emerge through adaptation of learning parameters then affecting each association individually. The second family included hybrid models derived from the Probe model [5], that explicitly handles multiple strategies built by Q-learning. Thus, although the Probe family is still based on incremental learning, it accounts for memory and generalization effects by directly representing and manipulating strategies. This family also included variants with learning rate adaptive control. The third family consisted of pure inference models, based on a complete set of possible strategies for the task. Hence, these models have access to a complete repertoire of strategies, which comprises the 3^3^ = 27 possible monkey-fruit mappings in our task, plus a completely random strategy.

**Figure 3:**
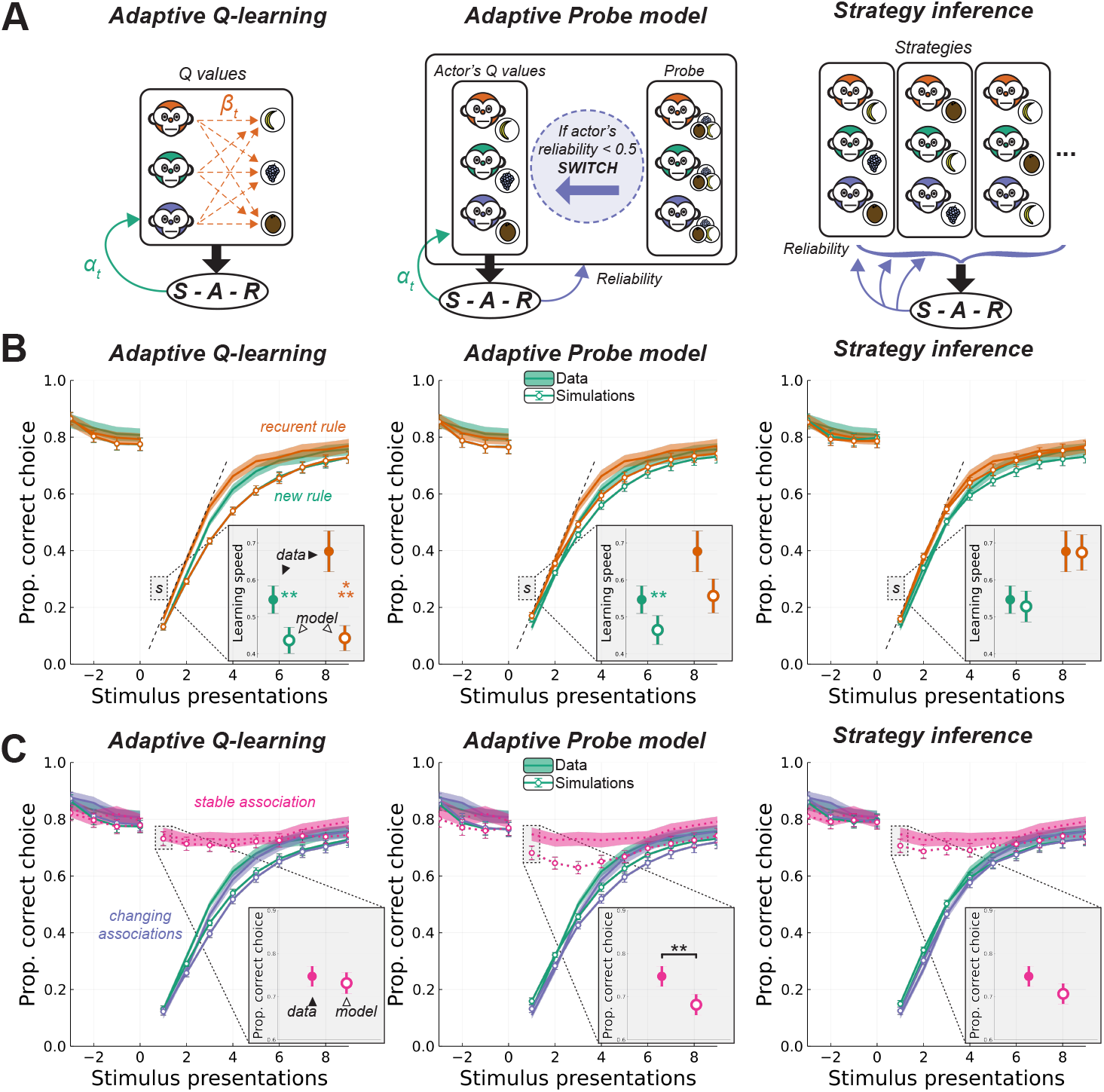
Description of model families and simulations: **A:** Diagrams of the different model families. **B & C :** Data (ribbons) and simulations (lines) of the main behavioral results (population mean ± standard error). Left column: adaptive Q-learning models could not account for the recurrence effect since it lacked a memory of past strategies, and did not reach similar learning speeds as human participants (insert: learning speed computed by fitting a saturating exponential, population mean± standard error). However the interference effect was well captured by the dynamic adaptation of learning parameters (insert: proportion of correct responses in the first presentation of the stable association, population mean ± standard error). Middle column: adaptive Probe models better captured the recurrence effect but overestimated the interference effect, with overall performance still inferior to that of human participants. Right column: Strategic inference models reproduced the overall performance of human participants, and captured well both recurrence and interference effects. ***: p < 0.001, **: p < 0.01, n.s.: non significant.

Instead of updating Q-values as in previous models, the Strategy Inference model computes them by marginalizing the reliability of each strategy. Reliabilities are updated according to Bayes rule for each strategy given the previous observation, controlled by a free parameter *ρ* corresponding to the evidence weight (the higher the value, the greater the impact of an observation on the update).

Crucially, the model also tracks a long-term prior over strategies simply by computing the average of the reliability over the whole experimental session. This long-term prior could therefore reflect the recurrence of certain strategies and account for the memory effect. Interestingly, the computational cost of this memory mechanism does not increase with the number of recurrent strategies (more recurrent strategies would simply share most of the probability mass in the long-term prior), which is consistent with the similarity of the behavioral results for versions of the task with one recurrent rule or three recurrent rules.

The Strategy Inference model has two variants. In the first variant (referred to as SI), all transitions between strategies are equiprobable, with the probability of changing strategy at each trial controlled by a fixed parameter *ω*. In other words *ω* corresponds to the perceived volatility in the task, with higher values corresponding to higher rates of changes. In the second variant (referred to as SI-Struct), the perceived statistical structure of the environment was more finely captured with three different parameters, *ω*_0_, *ω*_1_, *ω*_2_, corresponding to transitions between strategies with 0, 1, or 2 similar associations respectively, and two additional parameters for switching in and out of the random strategy (*ω*_*ri*_ and *ω*_*ro*_ respectively). Higher values of these parameters would reflect higher probabilities for transitioning, e.g. higher *ω*_0_ meaning higher probability of 0-similarity strategy changes.

We fitted all models to the data of each participant and simulated behavior using the set of fitted parameters. Then we performed similar analyses on the simulated data as on the human participants. Figure 3B and C shows the simulated performance of the best model of each family, over the performance of the human participants (see Supplementary Figure S4 for all models). Only models of the SI family captured all behavioral features described above and performed closer to humans than any other models. Firstly, the slope of the learning curve after a complete rule change was significantly lower than human’s performance for adaptive Q-learning model (Figure 3B left panel; new rule: −0.088, *CI*_95_ = [−0.137, −0.0407], Wilcoxon signed rank test *Z* = 312, *p* = 0.001; recurrent rule: 0.159, *CI*_95_ = [−0.273, −0.085], *Z* = 273, *p* < 0.001) and adaptive Probe model when facing a new rule (Figure 3B middle panel; new rule: −0.052, *CI*_95_ = [−0.108, −0.016], Wilcoxon signed rank test *Z* = 387, *p* = 0.009; recurrent rule: −0.079, *CI*_95_ = [−0.194, 0.003], *Z* = 464, *p* = 0.062).

In contrast, the learning slopes of simulated data from the SI-Struct model were not significantly different from humans’ (Figure 3B right panel; new rule: 0.008, *CI*_95_ = [−0.049, 0.078], Wilcoxon signed rank test *Z* = 715, *p* = 0.629; recurrent rule: −0.070, *CI*_95_ = [−0.083, 0.153], *Z* = 770, *p* = 0.318).

Secondly, the interference effect, measured as the difference in proportion of correct responses between the plateau and the first presentation of the stable stimulus after a partial rule change, was well captured by the adaptive Q-learning model (Figure 3C left panel; 0.025, *CI*_95_ = [−0.03, 0.07], Wilcoxon signed rank test *Z* = 701, *p* = 0.381) and the SI-Struct model (Figure 3C right panel; 0.04, *CI*_95_ = [−0.008, 0.098], Wilcoxon signed rank test *Z* = 812.5, *p* = 0.092), but significantly different from human performance with the adaptive Probe model (Figure 3C middle panel; 0.08, *CI*_95_ = [0.025, 0.1175], Wilcoxon signed rank test *Z* = 909, *p* = 0.009). More specifically, this model largely overestimates the interference effect for several trials after the rule change. This might be due to the fact that erroneous strategy change (leading to the interference) is more costly in the Probe model, since limiting the number of monitored strategies can lead to the loss of the correct strategy and the need to learn it from scratch. This observation supports the hypothesis of a more conservative mechanism, either by simply lowering the inverse temperature (as in the adaptive Q-learning model) or by direct inference over the close set of possible strategies (as in SI models).

Finally, we conducted a 6-fold cross validation to quantitatively compared all models (see Material and Methods) in the context with mixed complete and partial rule changes in the main sample. This context was ideal for model disambiguation since it was the only one with all the rule types (recurrent, complete and partial). This model comparison favored the SI-Struct model with an exceedance probability of 1 (Figure 4).

**Figure 4:**
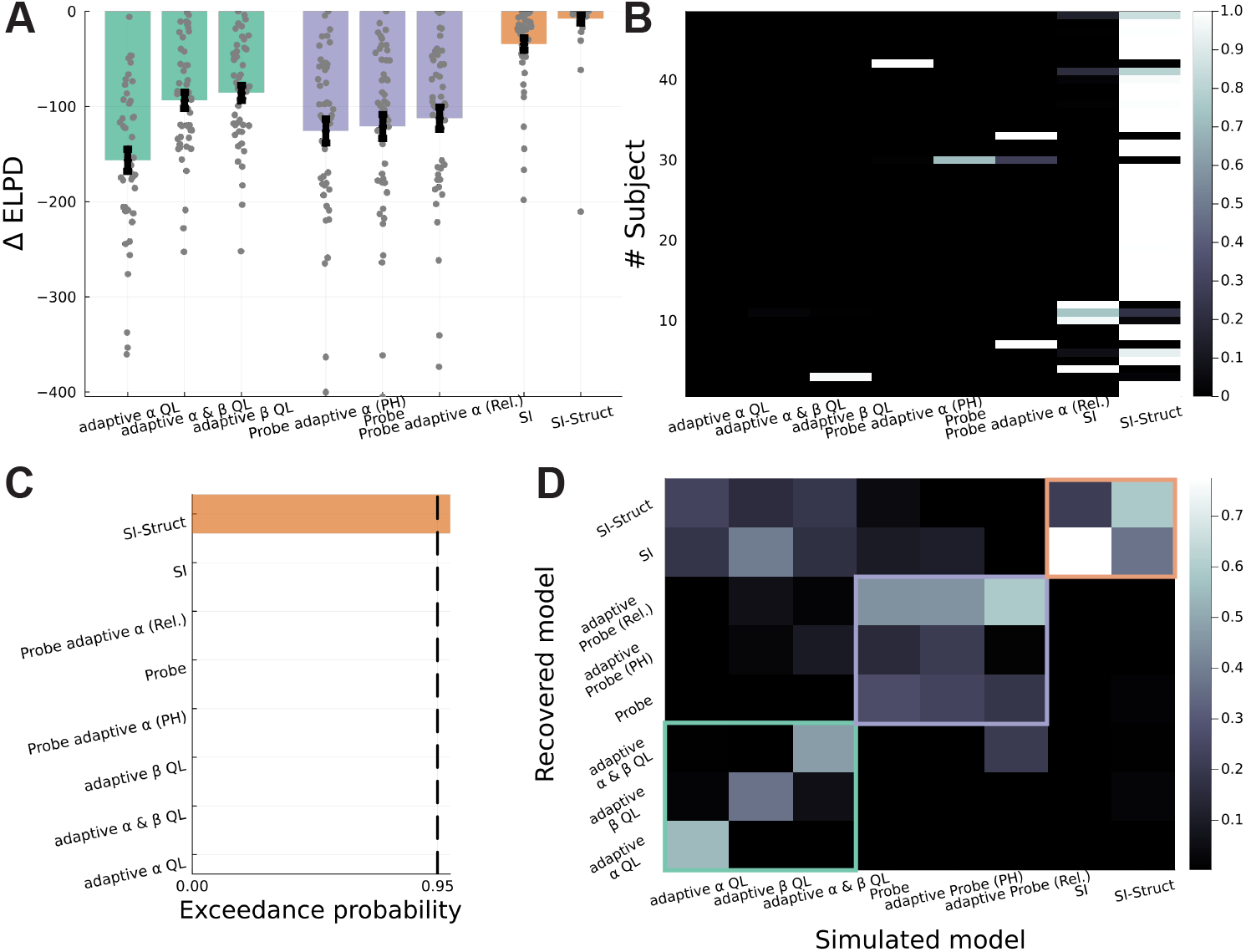
Quantitative model comparison: **A:** Distribution of the difference of expected logpredictive density (ELPD) compared to the best model across participants. The ELPD was computed from 6-fold cross validation. The closer to zero, the better. **B:** Attribution matrix : each row represents the probability of model attribution for a given participant. **C:** Exceedance probability of each model. **D:** Confusion matrix. While model families are quite distinct, there was a fair amount of confusion within families (colored squares).

### Contextual variation reveal a strong influence of the environment’s latent structure on strategy inference

In order to further investigate the reach and flexibility of strategy inference, we designed the two experimental contexts administered to the main sample of participants such that they have one identical type of rule change: partial rule changes with one stable association. This rule change was either alternating with complete rule changes in one context, or with partial rule changes with two stable associations in the other context. Thus, comparing the behavior of the same participants after 1-stable association partial rule changes between contexts would reveal a within-subject effect of the overall context.

Indeed, as seen in Figures 5A and B, the performances of the same participants, facing the same rule change (1-stable association) revealed a striking influence of the context. In the context with complete changes, participants learned changing associations faster (Figure 5A; difference in proportion of correct responses between contexts; significant cluster from presentation 2 to 5 after rule change; cluster-based permutation corrected *p* < 0.001), but showed a greater interference effect for the stable association (Figure 5B and insert). In the context without complete changes, the onset of the interference effect was delayed with no statistical difference on the first presentation of the stable stimulus compared to the plateau (median difference from plateau : 0.0, *CI*_95_ = [−0.054, 0.009], Wilcoxon signed rank test *Z* = 483.5, *p* = 0.201), though this difference was significantly different from the context with complete changes (median difference between context without complete changes and context with complete changes: −0.050, *CI*_95_ = [−0.087, −0.008] Wilcoxon signed rank test *Z* = 409, *p* = 0.018). This result is particularly remarkable since the same participants adapted differently to the same rule change depending on the latent structure of the environment without any indication of this variation.

**Figure 5:**
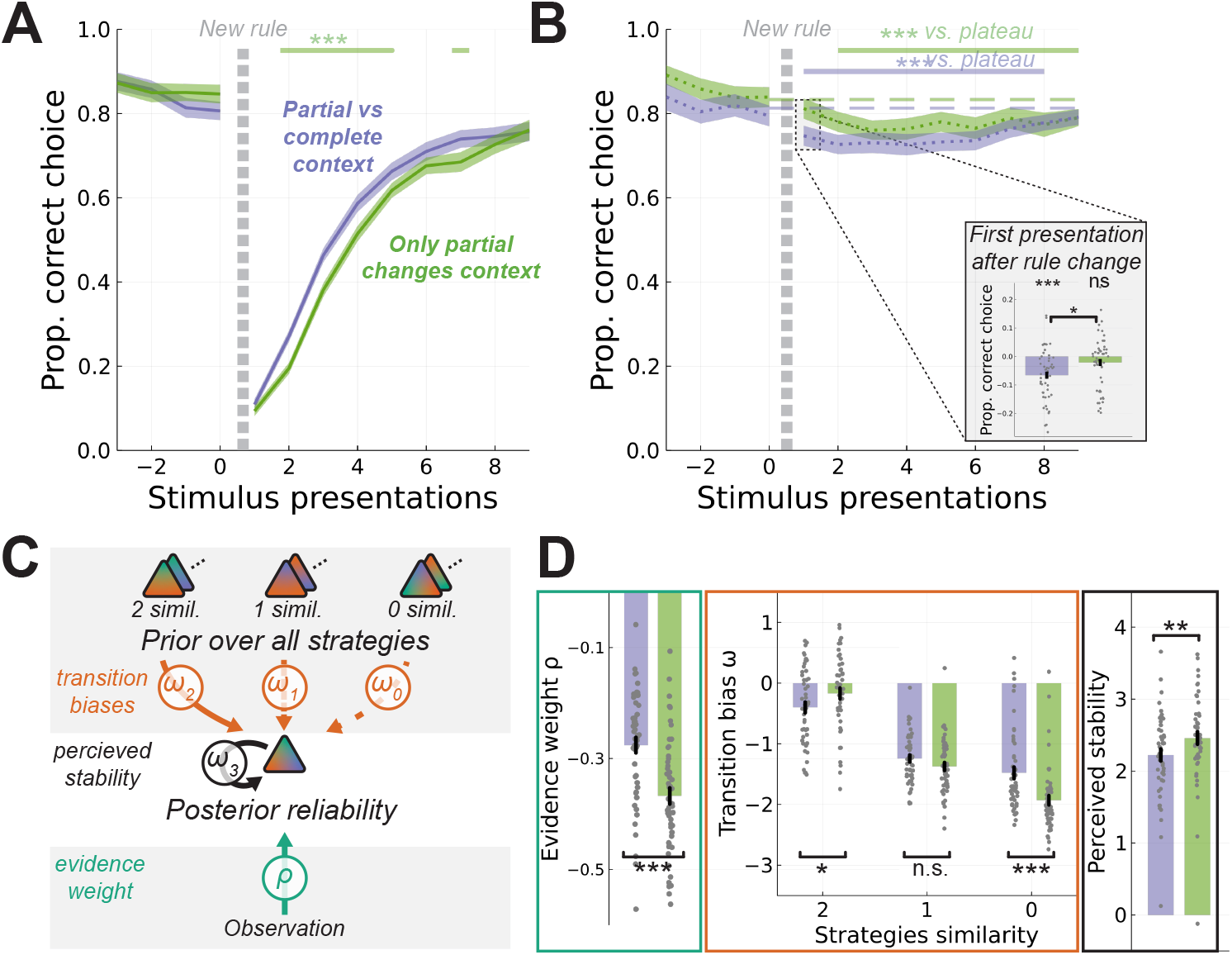
Strategy inference is modulated by latent contextual statistics: Comparison between similar rule changes in two different contexts, for the same group of participants. **A-B:** Behavioral performance after 1-stable association partial rule changes in both contexts (mean ± standard error). Ribbons on the top represent statistically significant effects after correcting for multiples comparisons with a cluster based permutation test. Participants learned faster and displayed a more pronounced global interference in the environment where rule changes were mostly complete (blue) compared to the environment when all rule changes were partial (green). **C:** Diagram of the SI-Struct model highlighting the parameters modulated by latent contextual statistics. Specifically, transition biases and perceived stability explicitly reflect the structure of the environment. Note, however, that this structure was never instructed to the participants. **D:** Values of fitted parameters of the SI-Struct model in both contexts. In the context without complete rule changes, the value of 4 parameters was significantly different than in the context with complete rule changes: the evidence weight (left), the transition probability towards 0-similarity and 2-similarities strategies (middle) and the perceived stability (right). ***: p < 0.001, **: p < 0.01, *: p < 0.05, n.s.: non significant.

Comparison of free parameter values of the SI-Struct model fitted separately to the data of both contexts revealed significant differences in specific parameters (Figure 5C and D and Supplementary Table 1): notably the evidence weight (*ρ*), the transition probability towards 0 and 2-similarities strategies (*ω*_0_ and *ω*_2_ respectively) and the perceived stability (*ω*_3_). Interestingly, these differences corresponded to real but hidden statistical differences between contexts that participants correctly inferred. Indeed, lower values for *ω*_0_ and higher values for *ω*_2_ indicate lower perceived probability to transition to dissimilar strategies, and higher perceived probability to transition to strategies with 2 similarities, which corresponds exactly to the difference in the type of rule changes between contexts.

**Table 1:**
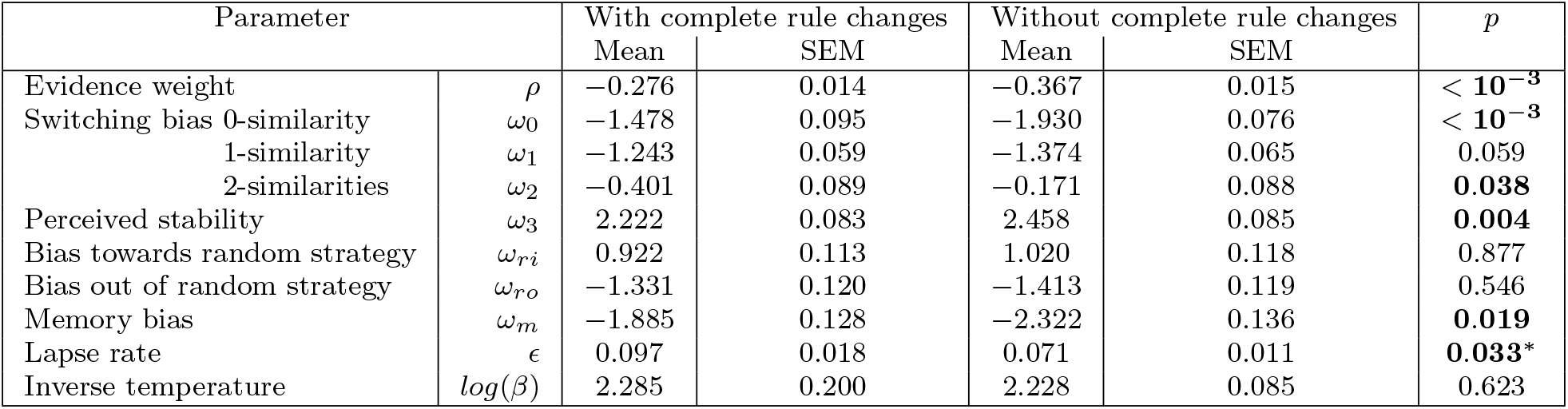
Fitted parameters of the Strategy Inference model in both contexts. SEM : Standard Error of the Mean. p : p-value for the Wilcoxon Signed Rank test comparing the values in context 1 and 2 for each participant. ^∗^When removing one outlier with an ϵ value of 0.696 in context 1, the difference was no longer significant (p = 0.052).

Moreover, the participants behaved in this second context as if individual observations were less informative (lower evidence weight *ρ*) and overall rule changes less frequent (higher perceived stability (*ω*_3_). This could be explained by several factors. First, in order to check the pervasiveness of global behavioral changes, feedback’s reliability for two stable associations was transiently lowered to 75% during twenty trials around half of 2-stable associations rule changes (9 over 18) and 9 more times randomly during the session without any rule change in the second context. While this is unlikely to have produced long lasting effects in the participants’ inference processes (two third of the trials over 20 trials episodes), it might have led to a decrease of the evidence weight. Second, for the same number of rule changes in both contexts, local associations remained the same for longer periods in the context with no complete rule changes than in the context with complete rule changes. Thus the frequency of local rule changes was lower in this second context, which could confound the effect of the task structure per se on the perceived volatility and/or evidence weight.

In order to rule out these potential confounding factors, we conducted the same analysis in the independent additional sample of participants. Crucially, for these participants the local rate of changes was matched in all three contexts: whether changes were complete or partial, each local rule for a stimulus was changed every 14.5 trials on average. This allowed to control for local volatility as a possible confounding factor for contextual modulation, and should thus less likely produce changes in the model’s evidence weight between context. As shown in Figure 6A, fitting the SI-Struct model to the data of the 3 contexts recovered a specific adaptation of inference parameters to the task structure (see Supplementary Table 2). Indeed, in this dataset, contextual differences impacted only the values of parameters controlling the transition probabilities between strategies, and not the evidence weight or any other free parameter. As in the previous experiment, these differences in parameter values reflected real differences in latent contextual statistics, even though participants were never given instructions on the structure of the rule changes and were simply informed that they had to redo the same task three times.

**Table 2:**
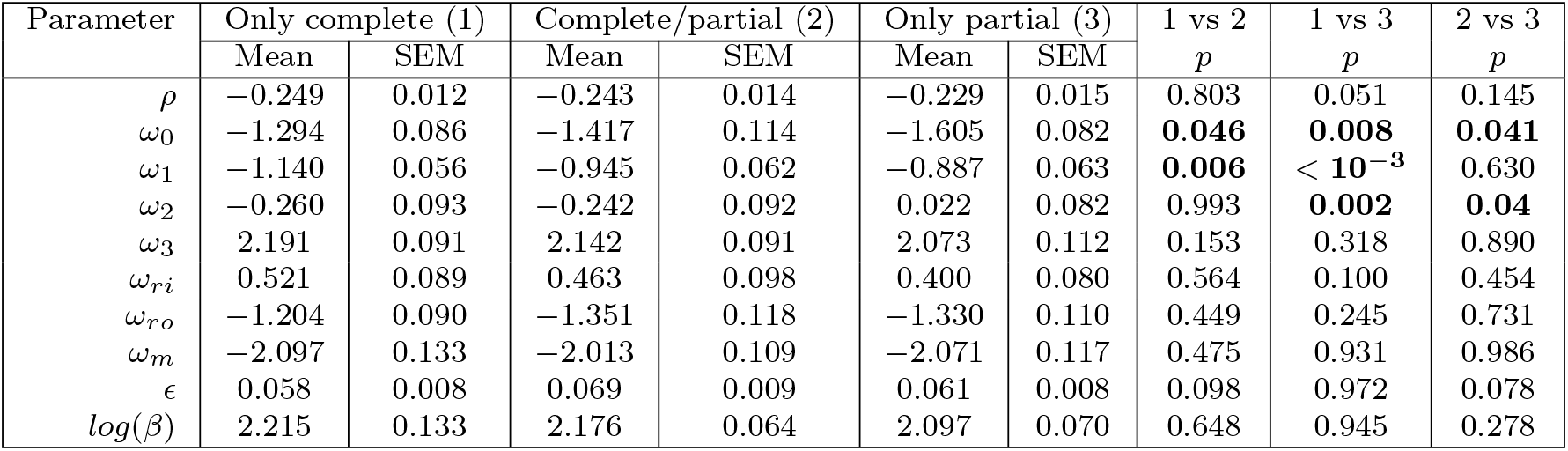
Fitted parameters values of the Strategy Inference model in the replication dataset. SEM : Standard Error of the Mean. p : p-value for the Wilcoxon Signed Rank test comparing the values in context 1 and 2 for each participant.

| Parameter | Only complete (1) |  | Complete/partial (2) |  | Only partial (3) |  | 1 vs 2 | 1 vs 3 | 2 vs 3 |
| --- | --- | --- | --- | --- | --- | --- | --- | --- | --- |
| | Mean | SEM | Mean | SEM | Mean | SEM | $p$ | $p$ | $p$ |
| $\rho$ | -0.249 | 0.012 | -0.243 | 0.014 | -0.229 | 0.015 | 0.803 | 0.051 | 0.145 |
| $\omega_0$ | -1.294 | 0.086 | -1.417 | 0.114 | -1.605 | 0.082 | **0.046** | **0.008** | **0.041** |
| $\omega_1$ | -1.140 | 0.056 | -0.945 | 0.062 | -0.887 | 0.063 | **0.006** | $< 10^{-3}$ | 0.630 |
| $\omega_2$ | -0.260 | 0.093 | -0.242 | 0.092 | 0.022 | 0.082 | 0.993 | **0.002** | **0.04** |
| $\omega_3$ | 2.191 | 0.091 | 2.142 | 0.091 | 2.073 | 0.112 | 0.153 | 0.318 | 0.890 |
| $\omega_{ri}$ | 0.521 | 0.089 | 0.463 | 0.098 | 0.400 | 0.080 | 0.564 | 0.100 | 0.454 |
| $\omega_{ro}$ | -1.204 | 0.090 | -1.351 | 0.118 | -1.330 | 0.110 | 0.449 | 0.245 | 0.731 |
| $\omega_m$ | -2.097 | 0.133 | -2.013 | 0.109 | -2.071 | 0.117 | 0.475 | 0.931 | 0.986 |
| $\epsilon$ | 0.058 | 0.008 | 0.069 | 0.009 | 0.061 | 0.008 | 0.098 | 0.972 | 0.078 |
| $\log(\beta)$ | 2.215 | 0.133 | 2.176 | 0.064 | 2.097 | 0.070 | 0.648 | 0.945 | 0.278 |

**Figure 6:**
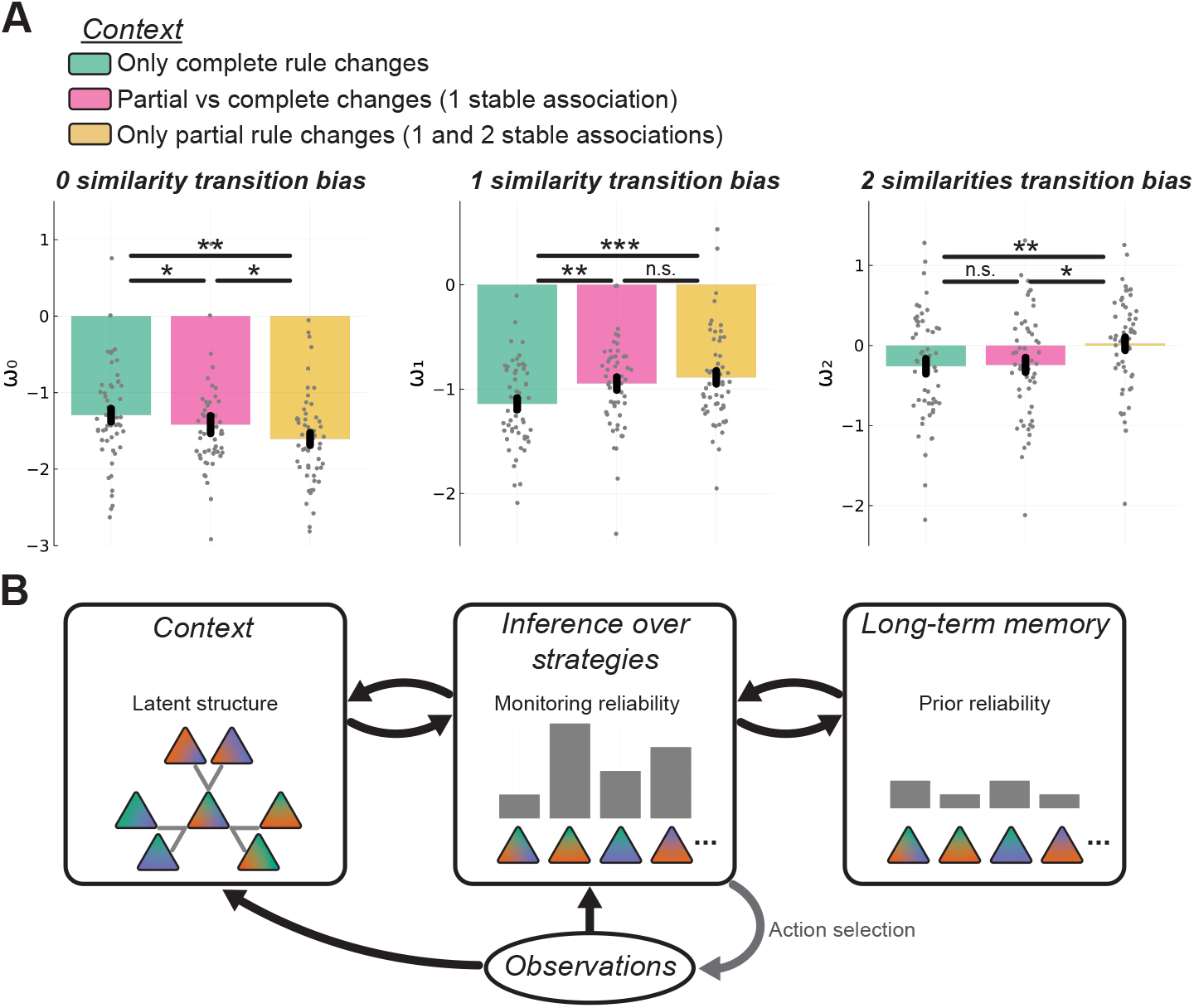
Inference within context. **A:** Influence of the contexts’ latent statistics on fitted parameters in the additional sample. Left: The fitted values of the perceived transition probability towards strategies with 0 similarity was significantly higher in the context with only complete rule changes compared to other contexts, and higher in the context with complete and partial rule changes compared to the context with only partial rule changes. Middle: The fitted values of the perceived transition probability towards strategies with 1 similarity was significantly lower in the context with only complete rule changes compared to other contexts. Right: The fitted values of the perceived transition probability towards strategies with 2 similarities was significantly higher in the context with only partial rule changes (including changes with 2 stable associations) compared to other contexts (without 2 stable associations rule changes). **B:** Diagram of the theoretical proposal. ***: < 0.001, **: p < 0.01, *: p < 0.05, n.s.: non significant.

## Discussion

In this study, we investigated the extent to which human participants can adapt to unexpected changes by inferring abstract strategies. To this end, we designed a novel task that directly tested a set of behavioral predictions regarding inference at a global strategy level, independent of the incremental learning of local stimulus-action associations. Not only did we find strong evidence that strategylevel inference is sufficient to account for the remarkable behavioral adaptability observed in the task, but we also demonstrated the high flexibility of strategylevel inference itself, as participants spontaneously adapted this process to the latent statistical structure of the environment.

First, we demonstrated a strategy-level memory effect in two independent groups of participants, revealing the rapid retrieval of previously used strategies from a working memory buffer (Figure 1C and Supplementary Figure S3). While this effect had been shown in a similar task [5], it had only been observed in a closed environment where three identical rules alternated. Here, we show that this effect is robust enough to persist in an open environment, where up to three recurrent rules are randomly interspersed with new rules. Moreover, this strategy-level memory effect could not be explained by memory or learning at the level of individual stimulus-action associations, as we found no performance differences between rare and frequent associations after rule changes (as shown in Supplementary Figure S2).

Second, we found that partial rule changes (i.e. restricted to some stimulus-action associations) interfered with the performance of the association that remained stable (Figure 1C). Using a Hidden Markov Model (HMM) to detect abrupt transitions in participants’ choice behavior without making mechanistic assumptions about how they were triggered, we showed that this interference effect specifically resulted either from erroneous inference at the strategy level (Figure 1D), which led participants to transiently use a non overlapping behavioral strategy (14.7 ± 1.9% of strategy changes after partial rule changes), or from switches to the random strategy (28.9 ± 2.5% of strategy changes after partial rule changes).

Third, these phases of seemingly random behavior were most likely bouts of exploration, during which participants actively probed their environment. Indeed, choices during these phases were not strictly equiprobable but were instead biased against the previously used strategy, suggesting an active search for information on alternative choices (Figure 2C). Moreover, switching out of the random strategy was abrupt, resulting in an instantaneous shift from below-chance performance to near-plateau performance (Figure 2D). Taken together, this suggests that participants engage in a random strategy to actively sample and accumulate information about their environment before switching to an exploitation regime once the correct strategy has been identified.

Decision models based on incremental learning of stimulus-action associations cannot account for these observations unless they incorporate additional computational mechanisms to enhance their flexibility. This can be achieved, for instance, by continuously adjusting learning and/or decision parameters [28, 31, 32] and/or by introducing a form of strategy-level representation. The latter approach involves tracking the reliability of previously encountered strategies, as in the PROBE model [5]; see also [9, 33, 34]). While these two broad categories of adaptation mechanisms – often referred to as meta-reinforcement learning and multi-modular models respectively – differ in terms of computational architecture, they are not mutually exclusive. In fact they can be integrated within the same models as illustrated by the adaptive PROBE model assessed in our study (see Figure 3) because both ultimately rely on a common adaptation signal: the local prediction error computed between a stimulus-action association and its outcome.

Local prediction error is central to our understanding of learning because of how successful it has been at explaining midbrain dopamine phasic activity, both biologically and theoretically [35, 36]. However, more recent studies show that midbrain dopamine activity also encodes beliefs about environmental latent statistical structure, integrating abstract representations from medial prefrontal cortical regions [36, 37]. Current meta-reinforcement models account for interactions between beliefs and local prediction errors through various adaptation mechanisms. Meta-reinforcement learners typically integrate local prediction errors over time to adjust their learning rate [32] or inverse temperature [28], or use related signals to detect discrete change points [25] when resetting to a prior policy accelerate adaptation [38]. When temporal integration is global, i.e. spans all stimulus-action associations, pooled local deviations can induce global behavioral changes, leading to an interference effect (as shown in Figure 3C left panel). But none of these models can protect previously learned stimulus-action associations from being overwritten when adapting to a novel environment, which make them unable to capture strategy-level memory effects (Figure 3B left panel). This representational issue is independent of the specific algorithm used to adjust model parameters.

On the other hand, multi-modular models inherently support the parallel representation of alternative strategies, which can encompass qualitatively different representations of the environment [33, 34], or complementary behavioral policies in the discrete or continuous domain [5, 9]. In these models, local prediction errors update the reliability of concurrent strategies, a process linked to neural activity in the medial and lateral parts of the anterior prefrontal cortex [7, 29]. Strategy reliability estimates can serve either as a direct selection signal, with the most reliable strategy driving decision-making (as in the PROBE model), or as a weighting mechanism where a policy is inferred by averaging over the space of strategies [9]. Crucially, multi-modular models also use local prediction errors to locally adjust strategies via incremental learning, in addition to monitoring them using their updated reliability. While both processes are necessary in environments where the strategy space is open, novel and/or so large that it exceeds the human brain’s representational capabilities, it can slow down adaptability in closed and reasonably limited strategy spaces, which are common in our daily life. Indeed, building a cohesive behavioral strategy from scratch takes times. Moreover, the fluctuation of strategies themselves due to incremental learning may sometimes compromise the stability of reliability-based inference. This limitation is obvious in all PROBE model variants (see the adaptive PROBE model in Figure 3 middle panels and additional variants in Supplementary Figure S4), which successfully replicate both interference and strategy-level memory effects, but fail to match the speed of human learning.

With the strategy inference model, we take a more direct route: all complementary strategies are represented, and their reliabilities are updated not by a classical prediction error of the local Q-value, but by the likelihood of the observed stimulus-action-outcome association given each strategy. This transforms the question “*how much does each choice yield for each stimulus?*” into “*what combination of correct choices should I be using at the moment ?*”. In our task, this modeling approach offers a more parsimonious account of human behavior than any model based on incremental learning (as shown in Figure 3, right panels, Figure 4, and Supplementary FigureS4). Taken together, our results demonstrate that, in our task, human adaptation to rule changes mostly relied on inference over strategy-level representations (e.g. strategy reliability) rather than gradual learning of local stimulus-action associations (e.g. Q-values). Similar results have been reported in a study with 4×4 stimulus-action associations and deterministic feedback [24]. The authors reported mixed results, showing both very rapid learning by human participants, well beyond the performance of a Q-learning model, but also very high plateau performance, this time captured only by Q-learning. Interestingly, the model proposed to account for the learning phase was initialized with a set of deterministic strategies progressively eliminated as feedback was collected. Our model generalizes this approach to probabilistic and dynamically changing environments and captures all the phases of our participants’ behavior in the task.

Finally, we demonstrate that the inference of global strategies depended on the perceived structure of the environment. In our model, participant’s beliefs about this structure were captured by free parameters representing transition bias between strategies based on their similarity. Remarkably, even in the absence of explicit cues regarding the actual structure of the environment or its variations, estimated transition biases reflected participant’s adaptation to the underlying structure of the task. These results show that inference over strategies is not simply a change point detection process, but is likely subordinated to high-level abstract representations, that are only captured by our top-down strategy inference model. Indeed, in our model, prior beliefs about strategies during the task is influenced by (1) a dynamic long-term prior capturing memory of recurrent strategies, and (2) an abstract map of strategies capturing the perceived stability and transition biases between similar strategies (see Figure 6B).

A growing body of literature illustrates how cognitive maps are implemented in the brains of several species, involving the entorhinal cortex and hippocampus, alongside anterior prefrontal cortical areas [39, 40]. Cognitive maps are believed to serve as the foundation for navigation in high dimensional abstract spaces and to enable generalization across domains [21, 22, 39, 41, 42, 43, 44]. In our task, such high-level manipulations of abstract concepts may also support the creation of a structured space of putative strategies even before engaging in the task. Convergent work has demonstrated a critical interaction between the hippocampus and subcortical regions for learning and memorizing behavioral strategies [45, 46, 47]. The medial prefrontal cortex has been suggested to play a central role either as monitoring each strategy reliability [7, 29] or as encoding previously learned strategies, reactivated during hippocampal replay [44, 46]. On the other hand, several authors have proposed to re-characterize the functions of prefrontal regions usually involved in learning in terms of cognitive mapping, based on the impressive flexibility of these areas to encode context-relevant variables [7, 42, 48, 49]. Hence, the explicit representation of task-dependent strategies at the algorithmic level, as in our work, could serve as a basis to study how flexible strategy inference might be implemented at the neural level.

## Acknowledgements

We thank Nils Kolling for his insightful and detailed comments on the article and Stefano Palminteri for his careful proof-reading. This work was supported by an ANR JCJC grant (ANR-21-CE37-0004) awarded to Ph.D., and doctoral grants awarded to S.B from LABEX BioPsy program and the “Defense Innovation Agency” (U17JRAP016).

## Author Contributions Statement

S.B and Ph.D. designed all the experiments, S.B. carried out data recording and all statistical analysis. All the authors contributed to the design of the computational models and the writing of the manuscript.

## Material and Methods

### Participants

The study was approved by Sorbonne University’s ethics committee (protocol no. CER-2020-85). All participants were recruited via e-mail and gave their informed consent before participation in the study. participants were screened for the absence of any history of neurological and psychiatric disease or any current psychiatric medication. On the first day of the study, participants were sent a link valid during two weeks to perform the tasks online.

#### Main sample

58 participants connected at least once, and 51 participants (31 female, age 33 ± 15) completed the 4 sessions and were included in the analysis. They were paid a fixed amount of 40€ plus a maximum bonus of 20€ proportional to their overall performance (mean: 15 ± 4.7€).

#### Additional sample

74 participants connected at least once, and 54 (42 female, age 29 ± 13) completed the 3 sessions and were included in the analysis. They were paid a fixed amount of 30€ plus a maximum bonus of 15€ proportional to their overall performance (mean: 8 *±* 5.6€).

### Experimental tasks

The tasks were programmed in JavaScript using the jsPsych framework, and all the scripts, data and links were supported by JATOS. The main study was hosted on MindProbe.eu, the server of the European Society for Cognitive Psychology (ESCoP), while the replication study was hosted on the local Brain Institute (ICM) server in Paris.

#### Main sample

On each trial, participants were asked to select one of the three fruits (grapes, banana or coconut) to feed one of the three monkeys (green, blue and orange) currently displayed on the screen. After fruit selection, a feedback screen appeared to indicate if the answer was correct (the monkey liked the fruit) or incorrect (the monkey did not like the fruit). Crucially, in order to maintain some degree of uncertainty on the latent rules underlying the task, this feedback was stochastic: in 90% of the trials, a monkey would like only its preferred fruit (that the participants had to learn) and dislike the two other fruits, while the remaining 10% of the trials consisted in trap trials where monkeys disliked their favorite fruit and liked the two others. The feedback gave no information regarding the non-chosen fruits.

Participants were informed that each monkey had only one preferred fruit at a given time, but one fruit could be preferred by at most 2 monkeys. Participants were instructed that each monkey would occasionally change their preferred fruit without any cue. There was no instruction on the structure of such changes (i.e. participants were not biased to believe that all 3 monkeys would change their preferred fruit at the same time).

Each participant had to complete 4 sessions of 39 episodes. Each episode lasted 20 to 60 trials. When participants reached a performance criterion, the episode lasted 10 trials more before ending. The performance criterion was to get 4 correct answers out of the last 5 trials and 2 consecutive correct answers for each monkey. The rule change for the new episode was one of 4 possible types, depending on the latent structure of the session (context):

#### Context with complete changes

1. **Complete new rule** (12 episodes): All the monkeys changed their preferred fruit at the same time, forming a new mapping. Most of the time these mappings were never encountered before, but in order to solve the combinatorial problem, a few rules were considered new when not encountered for more than 8 episodes (i.e. more than approximately 160 trials).
2. **Partial new rule** (10 episodes): Two out of three monkeys changed their preferred fruit, forming a new mapping.
3. **Recurrent rule** (10 episodes): All the monkeys changed their preferred fruit, forming the same recurrent mapping all along the session.
4. **Rare rule** (4 episodes): All the monkeys changed their preferred fruit, forming a new mapping. Not only this mapping was never encountered before, but it was formed by associations that were unused for more than at leat 5 episodes (i.e. more than approximately 100 trials). Since no difference was found in the participants’ performance between this condition and the complete new rule condition, we merged them in the main analyses.

#### Context without complete changes

1. **1-stable association rule change** (9 episodes): Two out of three monkeys changed their preferred fruit. This condition is similar to the Partial new rule condition in Context 1.
2. **2-stable associations rule change** (9 episodes): Only one monkey changed its preferred fruit.
3. **2-stable associations rule change + noise** (9 episodes): One monkey changed its preferred fruit, but for the two others, the probability of trap trials (i.e. inaccurate feedback) increased to 25% from ten trials before the partial rule change to ten trials after.
4. **No rule change + noise** (9 episodes): None of the monkeys changed their preferred fruit, but the probability of trap trials increased to 25% for two monkeys, from ten trials before the partial rule change to ten trials after.

The 4 types of rule change were in random order, balanced within blocks of 13 episodes. At the end of a block, participants had a 2-minute break. The next block started with the previous block’s last rule so this was not considered a rule change.

The order of the 2 contexts was counterbalanced pseudo-randomly between participants: half of them did 2 sessions in the context with complete changes first, the other half started with 2 sessions in the context without complete changes.

On each trial, participants had 3 seconds to answer, after which the trial was considered invalid. After 5 consecutive invalid trials a warning appeared on the screen. After a second warning, the whole session ended and was considered invalid.

#### Additional sample

The instructions, the structure of trials and blocks, were similar to the main sample. However, in order to match the global frequency of local rule changes, the length of the entire session was kept constant at 1564 trials, and the duration of each episode varied depending on the condition. The total number of local changes for each context was 108. The participants had to perform 3 sessions, each with a different context, with different rule change types:

##### Context with only complete changes

1. **Complete new rule** (18 episodes - 39 to 45 trials each): All the monkeys changed their preferred fruit, forming a new mapping.
2. **Recurrent rule** (18 episodes - 39 to 45 trials each): All the monkeys changed their preferred fruit, forming one of 3 recurrent mappings.

##### Context with complete and partial changes

1. **Complete new rule** (20 episodes - 34 to 38 trials each).
2. **Partial (1-stable association) new rule** (24 episodes - 31 to 35 trials each): Two out of three monkeys changed their preferred fruit.

##### Context with only partial changes

1. **1-stable association new rule** (36 episodes): Two out of three monkeys changed their preferred fruit.
2. **2-stable associations new rule** (36 episodes): Only one monkey changed its preferred fruit.

(NB: In this context, since local changes were more intertwined, the number of trials per episode was such that one monkey would not change its preferred fruit twice in less than 30 trials. Thus, even if the global rule set often changed, local associations were maintained long enough for the task to be manageable.)

### Fitting the learning curves

In order to compute the learning speed presented in Figure 3, we fitted a function of the performance as the number of stimulus presentation from the rule changes:

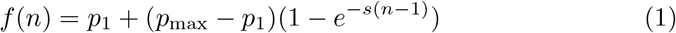

With 0 ≤ *p*_max_ ≤ 1 the asymptotic performance, 0 ≤ *p*_1_ ≤ 1 the performance for *n* = 1 and *s* ≥ 0 the learning slope. We let *s* vary with the condition, and *p*_1_ had different values for stable and changing associations. This model was fitted for each participant, with an adaptive Hamiltonian Monte Carlo scheme [50].

### Identification of global behavioral switches

In order to detect global behavioral switches from the dataset, we made the assumption that at any trial each participant would act according to a strategy, *i*.*e*., a stimulus-action mapping. Thus, we used a hidden Markov model (HMM), with the underlying strategy of the participant as the hidden variable *X*. Since there are 3 stimuli and 3 possible actions in our task, the set of all the possible strategies has a cardinality of 27. We added a 28th strategy corresponding to random choices for all 3 stimuli. Hence the vector **X** of all strategies is indexed from 0 to 27, with *X*_0_ the random strategy.

#### Parametrization of the HMM

We assumed a constant emission probability, i.e. the probability of acting according to the underlying strategy and not randomly, 0≤ *ρ* ≤ 1. The transition probabilities between strategies were parameterized by a vector **Θ** such that:

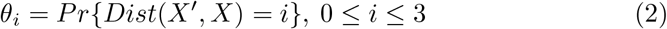

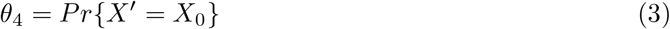

With *X*^*′*^ and *X* the next and current strategy respectively, and *Dist*(*X*^*′*^, *X*) the distance between strategies. The distance was simply the number of differences between the 2 mappings, hence *θ*_0_ corresponds to the probability of keeping the same strategy between 2 consecutive trials, while *θ*_3_ is the probability of a global behavioral switch. Finally, the free parameter 0 ≤ *τ* ≤ 1 controlled the probability of staying in the random strategy (hence 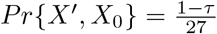 for all *X*^*′*^).

#### Fitting HMM parameters

In our HMM there were 7 free parameters, *ρ*, **Θ** = {*θ*_0_, *θ*_1_, *θ*_2_, *θ*_3_, *θ* _4_} and *τ*. We drew samples from their posterior distribution using Hamiltonian Monte Carlo sampling [50] for each experimental session and each participant. The log-likelihood of the parameters given the sequence of stimuli (*i*.*e*., monkeys) **S** and choices (*i*.*e*., fruits) **A** is:

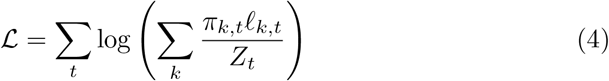

where *π*_*k,t*_ is the prior probability of the strategy *k* on trial *t*, 𝓁_*k,t*_ the likelihood for strategy *k* on trial *t* and *Z*_*t*_ the normalization constant. Thus:

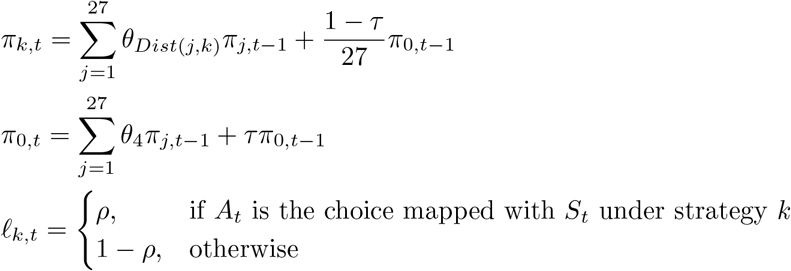

with 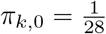 for all *k*.

### HMM inference and global switches detection

HMM inference refers to the process of inferring the most likely sequence of hidden variables given a sequence of observations and a set of parameters. With our samples from the parameters’ posterior, we could generate samples from the posterior over the hidden variables, i.e. the strategies used on a trial-by-trial basis, using Viterbi algorithm. Then, we were able to identify for every episode the trial with the highest probability of switch, and the identity of such a switch (switch to a random strategy, global strategic switch or overlapping strategic switch).

### Generative models

Three families of generative models were fitted to the data and simulated. In all models, actions (fruits) were sampled from a semi-uniform Boltzman (softmax) distribution:

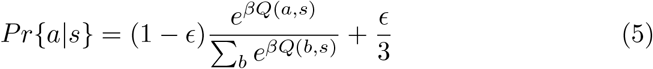

With *β* ≥ 0 the inverse temperature, and 0 ≤ *ϵ* ≤ 1 the lapse rate.

### Q-learning models

These models were variants of a basic counterfactual Q-learning model [51], in which Q-values were updated as:

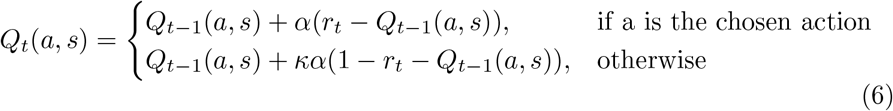

With 0≤ *α ≤* 1 the factual learning rate and 0 ≤ κ ≤ 1 the counterfactual learning rate. This parametrization insures that the counterfactual learning rate was always lower than the factual one, but followed the same dynamics (in the case of an adaptive learning rate).

#### Adaptive learning rate

The first variant featured an adaptive learning rate as in the classical Pearce-Hall model [32]:

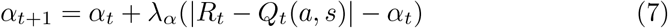

With 0 ≤ *λ*_*α*_ ≤ 1 a meta-learning rate.

#### Adaptive inverse temperature

In the second variant the inverse temperature *β* was adapted to the dynamics of the average reward rate, as in [28]:

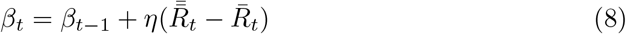

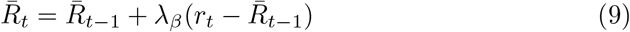

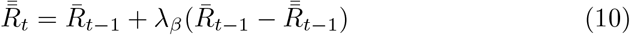

With 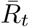 and 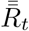 the first and second order moving averages of the reward rate respectively, *λ*_*β*_ a meta-learning rate and *η* ≥ 0 a scaling factor.

### Adaptive Probe models

These models were variants of the Probe model [5] where several sets of Q-values compete to be the actor strategy. Q-values are learned by reinforcement learning, and strategies are compared according to their reliability, *i*.*e*. the posterior probability of reflecting the actual rule conditioned by the history of previous observations. While several strategies can be monitored simultaneously, only the actor is updated by Q-learning. The size of this monitoring set is bounded, but the model is equipped with a long-term memory where previous actors are transferred when no longer relevant. When none of the monitored strategies predicts the current observations well enough, a probe strategy takes over from this long-term memory, until one of the monitored strategy or the probe can become the actor strategy. In this study, we have made a few modifications to the implementation of the original Probe model, to take into account the specific features of our experimental paradigm and to facilitate the fitting procedure:

1. Contrary to the original task [5], the same action (fruit) could be correct for more than one monkey, hence Q-values could not be normalized across stimuli. Instead, we used the same counterfactual-learning procedure as the Q-learning models (equation 6).
2. We removed the entropy-based computation of the probe’s initial reliability, in favor of a free parameter for probe initialization. We made this change for fitting convenience as we found that the original parametrization led to bad convergence.
3. We fitted the model with only one memory slot (in addition to the probe) for identifiability purpose, since, in the main study, there was only one recurrent rule and the long-term memory would already be biased by it. We checked that this would not disadvantage the model by fitting variants with 2 or 3 memory slots. As expected, adding extra memory slots did not improve the model fits.

#### Adaptive learning rate

Both variants of the Probe models had adaptive learning rates. The first one was based on the Pearce-Hall model described in equation 7. The other one used the actor’s reliability in order to compute the current learning rate:

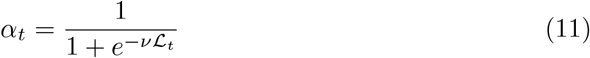

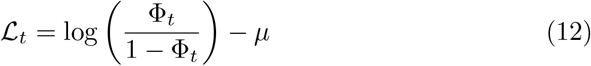

With Φ_*t*_ the reliability of the actor a trial *t. ν* and *μ* are free parameters defined on ℝ. Note that this parametrization did not impose a direction on the relation between the reliability and the learning rate. For positive values of *ν*, the learning rate increased with the reliability of the actor strategy, while it decreased for negative values.

### Strategic inference models

In these models, Q-values were not updated directly but computed by marginalizing the reliability of pre-defined strategies. On each trial, the Strategy Inference model computes the Q-value of each monkey-fruit association by marginalizing the reliability Φ of strategies that include that association:

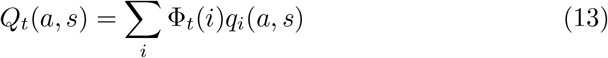

Where *s* (*stimulus*) denotes a particular monkey, *a* (*action*) denotes a particular fruit, and *q*_*i*_(*a, s*) = 1 if the strategy *i* includes the association {*a*| *s*}, and 0 otherwise.

The reliability Φ corresponds to the posterior probability that a given strategy reflects the current task rule, and is updated according to the Bayes rule:

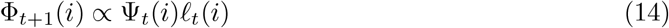

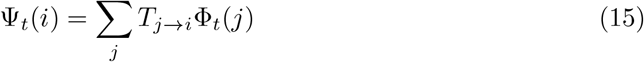

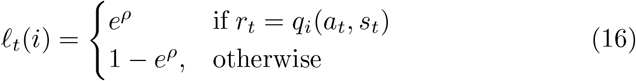

With Ψ(*i*) the prior of strategy *i, T*_*j*→*i*_ the transition probability from strategy *j* to strategy *i*, and 𝓁(*i*) the likelihood of the feedback *r* given the monkeyfruit pair {*a, s*}. This likelihood, *i*.*e*. the probability of the observed feedback *r* under a given strategy *i* was controlled by a free parameter *ρ* as in equation 16.

The model had two variants depending on the parametrization of the transition probability *T*. In the first variant (referred to as *SI*), all transitions were equiprobable, with the probability of changing strategy controlled by a fixed parameter *ω*. In the second variant (referred to as *SI-Struct*), *T*_*j*→*i*_ was controlled by different parameters depending on the similarity between strategies *i* and *j*. Hence this second variant captured the participant belief regarding the statistical structure of the environment (session) she/he is working in. Three different parameters, *ω*_0_, *ω*_1_, *ω*_2_, corresponded to transitions between strategies with 0, 1 or 2 similar associations respectively, and two additional parameters for switching in and out of the random strategy (*ω*_*ri*_ and *ω*_*ro*_ respectively).

To account for the memory of previously used strategies, the model included a long-term prior Γ over the distribution of strategies, updated via a moving average of the trial-by-trial posterior [38]:

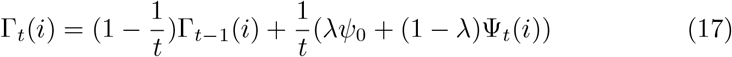

With *λ* a forgetting factor [38] and *ψ*_0_ a fixed prior (which was set to 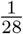) acting as an attractor when the forgetting factor was close to 1. Note the hyperbolic decay of the update, to counter recency effects and to allow the long-term prior to reflect the distribution of strategies over the entire session.

Hence, for each pair of strategies *i, j* with similarity *x*, the transition probability was computed as:

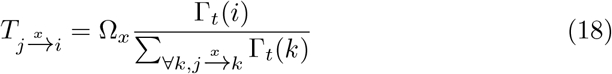

With 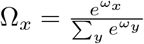. In the first variant, this simplifies to :

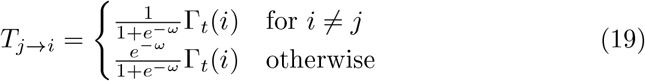

### Model fitting and model comparison

Models were fitted using the MCMC package Turing for Julia 1.6 [52]. Every model, except the Probe model, were fitted using an adaptive version of Hamiltonian Monte Carlo [50]. The Probe model could not, due to discontinuities in the likelihood function because of strategies selection, probe creation and deletion. Thus, we used a Gibbs sampling scheme, separating continuous parameters (learning rates, inverse temperature and lapse rate) that were fitted using Hamiltonian Monte Carlo, from discontinuous parameters (volatility, probe’s initial reliability and memory weight), that were fitted using a slice sampling scheme. For all models, convergence was assessed by qualitative inspection of the chains, and checks of the 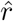 and *ESS* statistics.

For model comparison, we derived the predictive accuracy of each model via a 6-fold cross validation scheme, adapted for time-series [53]. Indeed, due to the temporal dependency of the latent variables, the models had to be fitted to all the data up to the testing block. For each participant, each of the two sessions was divided into 3 blocks of equal length that were used as testing datasets. The training datasets for each block comprised all the blocks of the other session, plus the previous blocks (if any) of the same session. We then derived the expected log-predictive density as defined in [54], which was used to compute attribution frequencies and exceedance probabilities [55].

## Supplementary material

**Figure S1:**
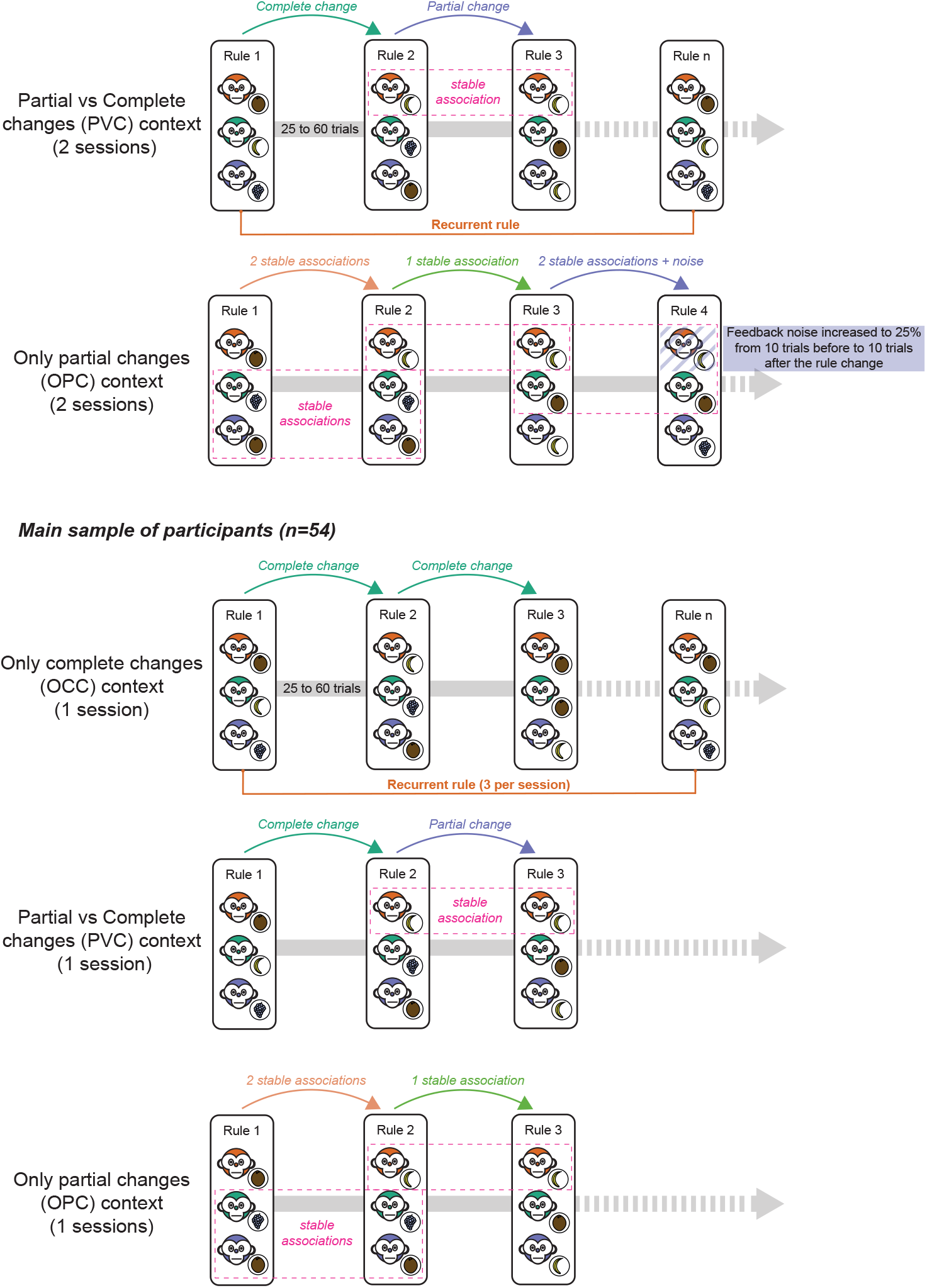
Diagram of experimental design. A main group of participants (upper panel) performed the task composed of 2 experimental contexts (2 sessions each). Fifty one participants completed all the sessions and were included in the analyses. An additional group of participants (lower panel) performed the task composed of 3 experimental contexts (1 session each). Fifty four participants completed all the sessions and were included in the analyses.

**Figure S2:**
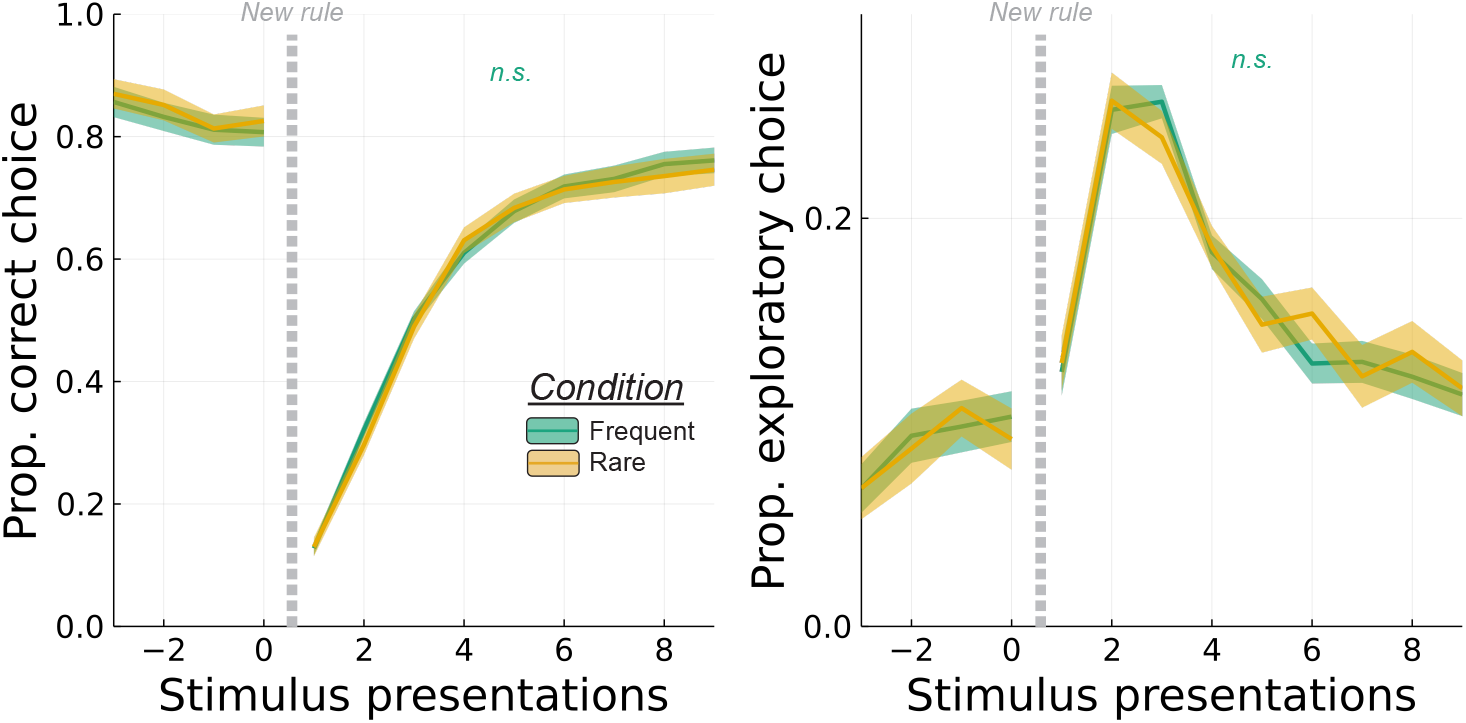
Testing local memory after complete rule changes: New rules could be composed from rare associations (which had not been observed in at least the last 5 episodes) or from frequent associations. There was no difference between these two conditions either in terms of correct choices (left panel) or exploratory choices (right panel).

**Figure S3:**
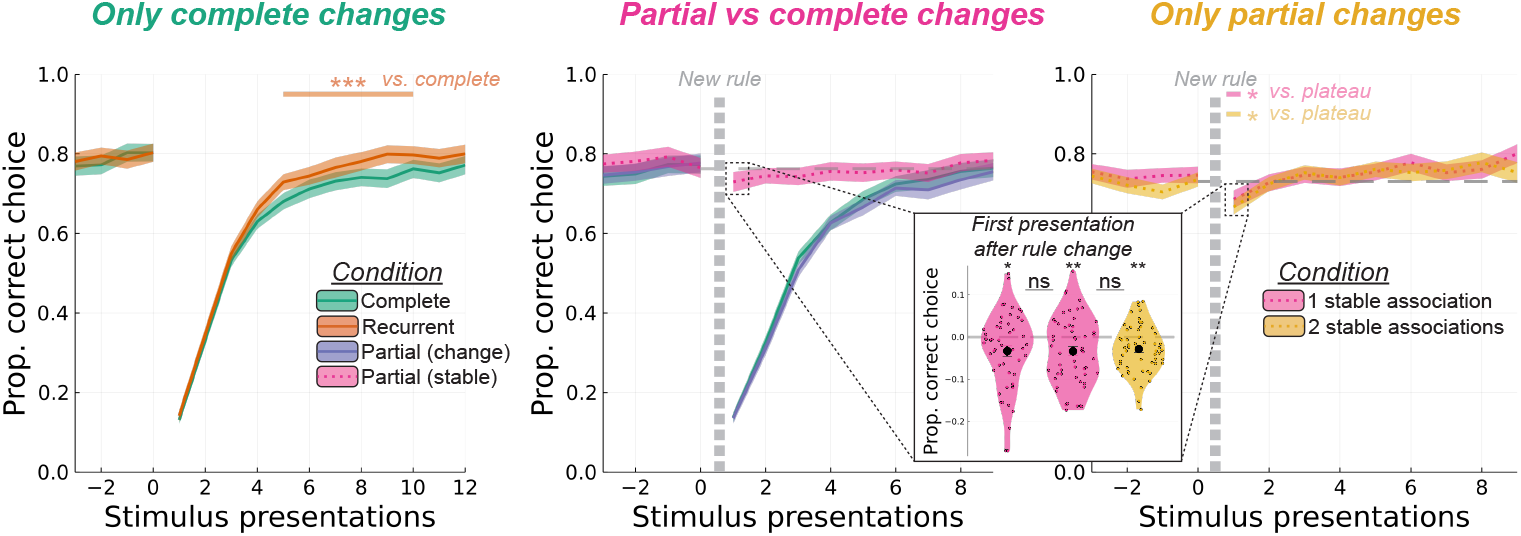
Behavioral results in the additional sample of participants. Ribbons on the top represent statistically significant effects after correcting for multiple comparison with a cluster based permutation test. **Left**: In the context with complete rule changes only and three recurrent rules alternating with random new rules, relearning was significantly faster when facing one of the recurrent rules compared to new rules. **Middle**: In the context with complete rule changes and 1-stable association partial rule changes, participants showed a significant interference on the first presentation of the stable association after rule change. Note that, although this effect is less pronounced than in the main task, it is still present with 24/44 rule changes being partial (10/36 in the main task). **Right**: In the context with partial rule changes only, the interference effect is still significant for both 1-stable association and 2-stable associations rule changes. Learning curves for the changing associations were not significantly different (data not shown). Insert: proportion of correct responses in the first presentation of the stable association, population mean ± standard error. ***: p < 0.001, **: p < 0.01 *: p < 0.05.

**Figure S4:**
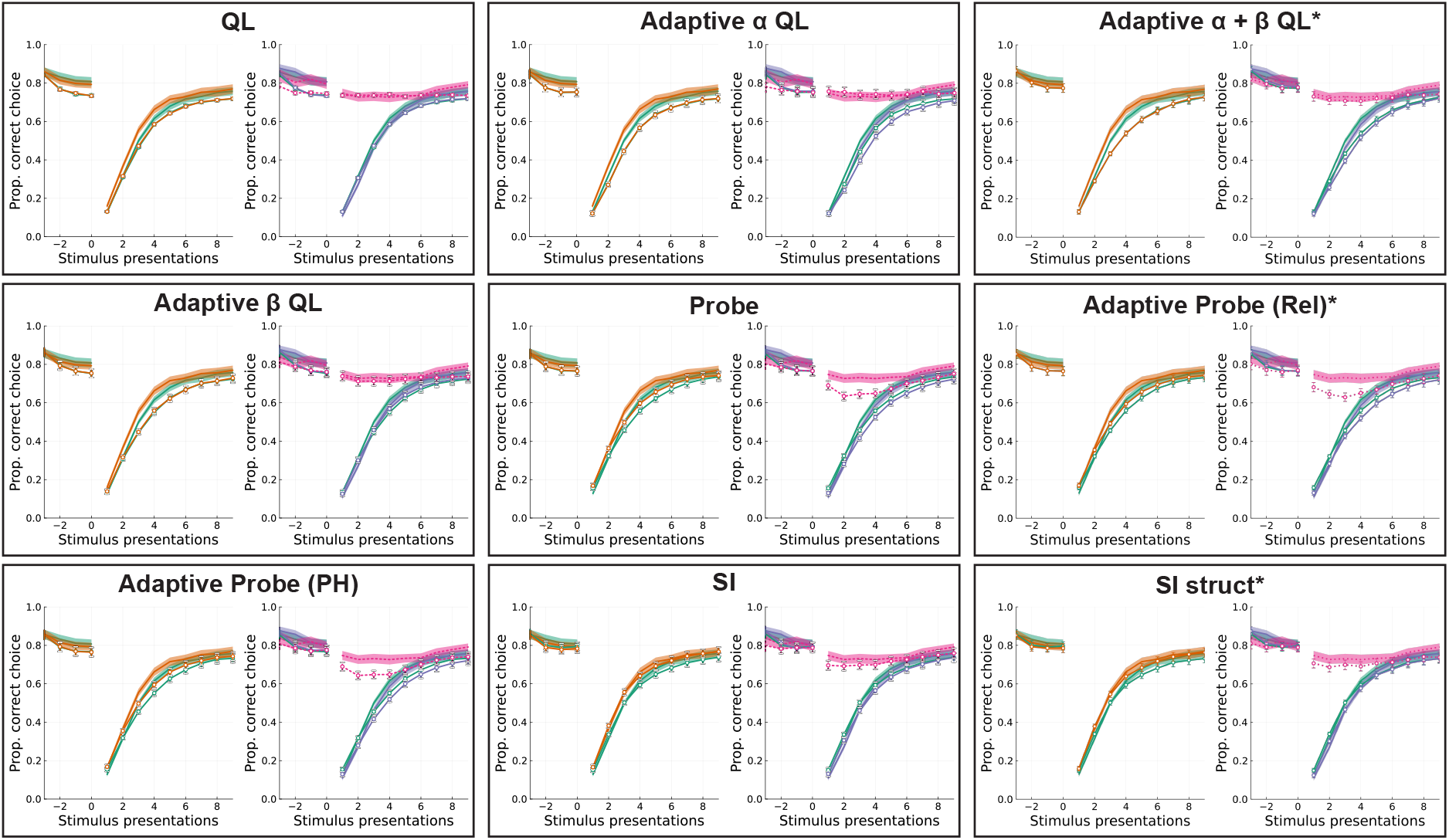
Simulations of all generative models: For each model, human data (ribbons) and simulations (lines) with fitted parameters. Left: proportion of correct responses after complete rule changes (green: new, orange: recurrent). Right: proportion of correct responses after partial rule changes (blue: changing association, pink: stable association) and new rule (green). Models with an asterisk after their name are winning models of their family.

**Figure S5:**
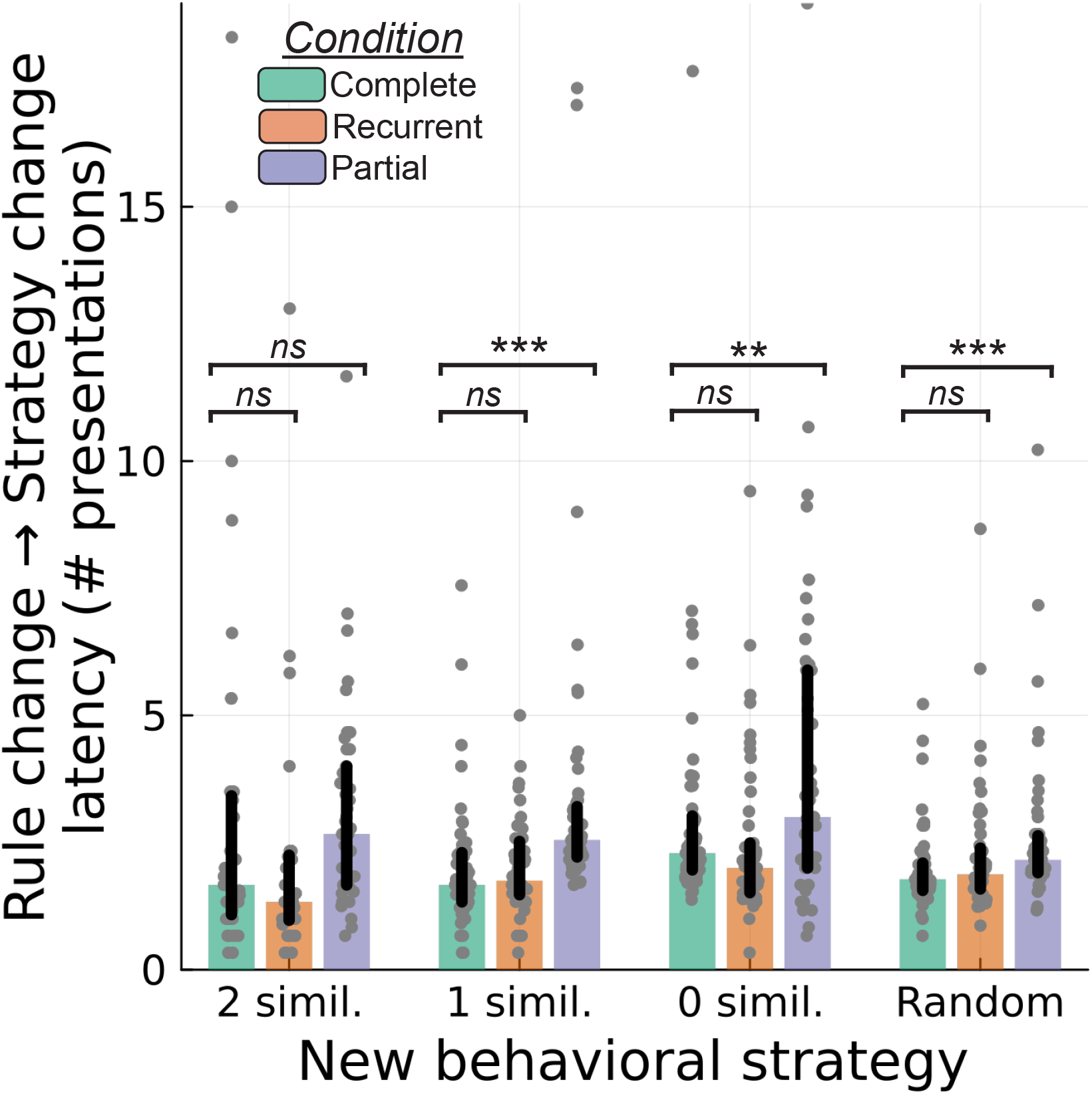
Rule change to strategy change latency: Number of presentations from the rule change to the strategy change (population mean ± standard error). All types of strategy changes were delayed after partial rule changes, compared to complete rule changes. ***: p < 0.001, **: p < 0.01.

**Figure S6:**
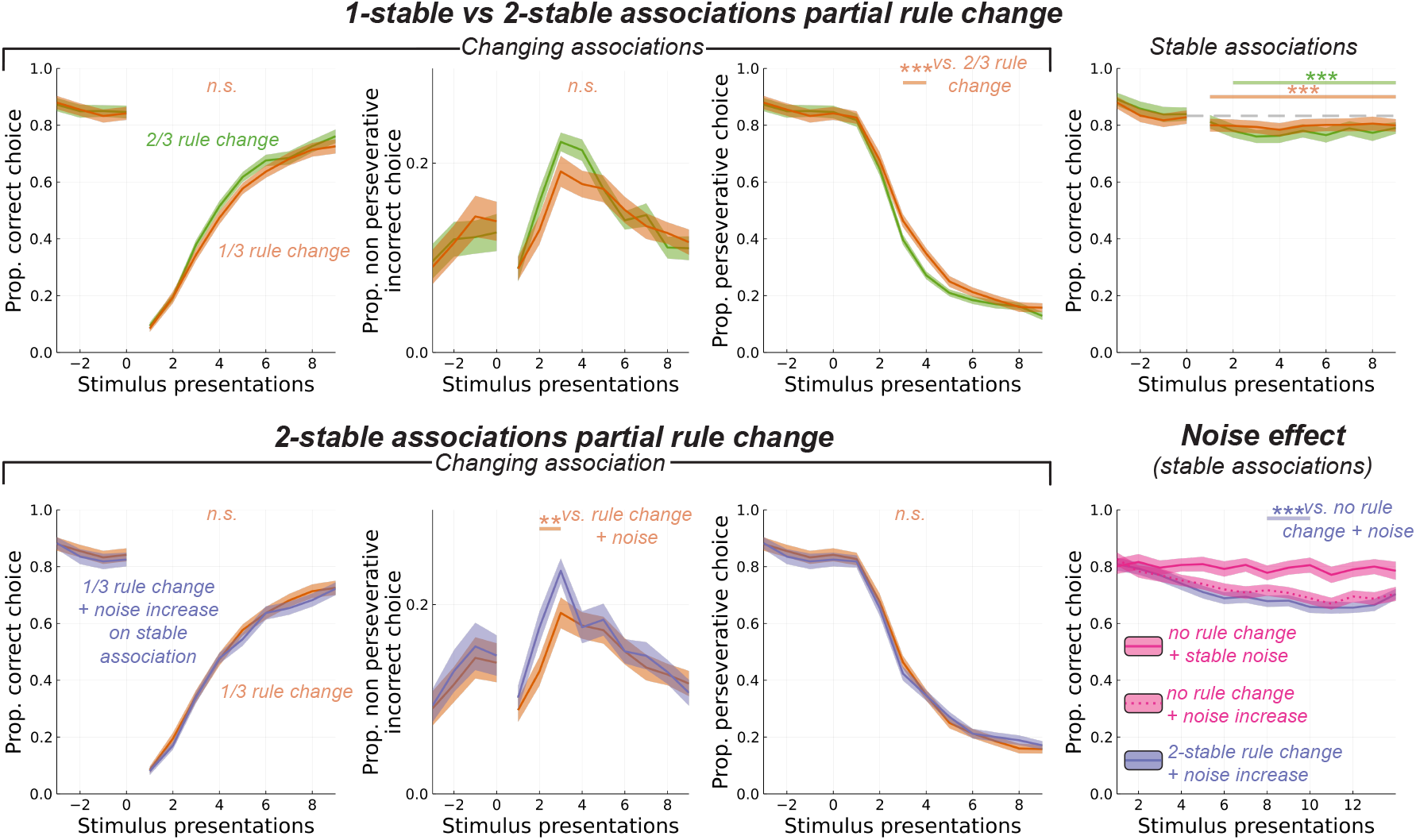
Behavioral results in the second environment. Ribbons on the top represent statistically significant effects after correcting for multiple comparison with a cluster based permutation test. **Upper pannel**: Comparison between 1-stable association rule change and 2-stable associations rule change conditions. Subjects persevere less after a 1-stable association rule change compared to a 2-stable associations rule change. They tend to explore more (non perseverative incorrect choices), though the difference is not statistically significant. Performance for stable associations is lower than the plateau for both conditions. **Lower pannel, left**: Comparison between 2-stable associations rule change with and without noise increase. The noise increase only affected stable associations. participants gave more exploratory answers for the changing association when stable associations were more noisy, though this effect did not significantly impact their performance. **Lower pannel, right**: Performance for the stable associations without noise increase or rule change (pink, solid line), with noise increase but no rule change (pink, dotted line) and with noise increase and rule change (blue, solid line). The interference due to rule change goes slightly beyond the effect of increasing noise (blue line vs pink dotted line). ***: p < 0.001, **: p < 0.01

